# Temporal hierarchy of intrinsic neural timescales converges with spatial core-periphery organization

**DOI:** 10.1101/2020.06.12.148866

**Authors:** Mehrshad Golesorkhi, Javier Gomez-Pilar, Shankar Tumati, Maia Fraser, Georg Northoff

## Abstract

The human cortex exhibits intrinsic neural timescales that shape a temporal hierarchy. Whether this temporal hierarchy follows the spatial hierarchy of its topography namely the core-periphery organization remains an open issue. Using Magnetoencephalography data, we investigate intrinsic neural timescales during rest and task states; we measure the autocorrelation window in short (ACW-50) and, introducing a novel variant, long (ACW-0) windows. We demonstrate longer ACW-50 and ACW-0 in networks located at the core compared to those at the periphery with rest and task states showing a high ACW correlation. Calculating rest-task differences, i.e., subtracting the shared core-periphery organization, reveals task-specific ACW changes in distinct networks. Finally, employing kernel density estimation, machine learning, and simulation, we demonstrate that ACW-0 exhibits better prediction in classifying a region’s time window as core or periphery. Overall, our findings provide fundamental insight into how the human cortex’s temporal hierarchy converges with its spatial core-periphery hierarchy.

## Introduction

The brain shows a complex temporal structure during both resting and task states. Recent findings demonstrate intrinsic neural timescales in resting state with shorter timescales in unimodal sensory regions and longer ones in transmodal regions of the default-mode (DMN) and fronto-parietal (FPN) networks ^1–4^. At the same time, there is increasing evidence that task states structure information in temporal terms as described by “temporal receptive fields” ^5^ or “temporal receptive windows” ^6^. Specifically, short segments like single words are processed mainly in unimodal lower-order sensory regions like the visual or auditory cortex while longer segments, integrating different stimuli, are processed in higher-order regions like DMN and FPN ^6–11^. How such temporal structuring during task states is related to the resting state’s intrinsic neural timescales remains yet unclear, though. One goal of our paper is, therefore, to investigate whether the resting state’s intrinsic neural timescales are preserved and/or changed during the transition from rest to task.

Analogous to the temporal hierarchy of intrinsic neural timescales, the hierarchical organization has also been observed on the spatial side, that is, its topographic organization. Converging evidence shows micro- and macro-scale hierarchical organization in the human cortex following what has been described as ‘core-periphery’ ^12–19^. A core-periphery (CP) architecture ^20^ can be characterized by a core that includes nodes with strong functional connectivity (FC) among each other (i.e. core-core connections or ‘rich club’ ^21–24^). These core-core connections can be observed among regions included in the default-mode (DMN), fronto-parietal (FPC), and cingulo opercular networks ^12–16,19^. These networks constituting the core have been distinguished from those at the opposite end of the core-periphery gradient, that is, sensory networks like visual, auditory, and somatomotor networks ^12–16,19^. The spatial hierarchy of CP has been mainly observed in functional magnetic resonance imaging (fMRI) measuring infraslow frequency ranges (0.01 to 0.1Hz). This leaves open whether CP organization is also present in the faster frequency (1.3 - 50Hz) as measured in MEG/EEG where, so far, an anterior-posterior hierarchy has been demonstrated ^25^. Therefore, our goal was to probe whether the core-periphery hierarchy is also present in the faster frequency range (1.3 - 50Hz) while controlling for anterior-posterior hierarchy.

How are temporal and spatial hierarchies related to each other? Recent studies demonstrate that the intrinsic neural timescales within specific regions strongly correlate with their FC ^1,3^. Given this relationship between FC and ACW, the FC-based spatial-topographic core-periphery organization may converge with the temporal hierarchy of the intrinsic neural timescales during both rest and task states. This touches upon the more general question about the relationship and potential convergence of temporal and spatial hierarchies in the brain, and whether such convergence is carried over from rest to task states. Investigating the convergence of spatial and temporal hierarchies of the brain is the overarching aim of our study. Insight into such possible convergent temporo-spatial core-periphery hierarchy is of importance to understand the brain’s organization which, in turn, is important for mental features like self and consciousness as they have neurally been characterized by temporo-spatial dynamics ^26–33^.

### Aims and hypotheses

The general aim of our study is to probe whether the temporal hierarchy of intrinsic neural timescales during both resting and task states follows the spatial topography of the core-periphery (CP) hierarchy (see Figure 1 for a general overview). To this end, we used MEG rest and task data from the Human Connectome Project’s (HCP) dataset. To operationalize core-periphery organization, we used three different definitions of CP including Schaeffer/Margulies (SCP) ^14^, Ji/Ito (JCP) ^1^, and restricted Ji/Ito (RCP). We determined the autocorrelation window (see below) of the regions/networks in the three different CP definitions to probe whether the hierarchical distribution of intrinsic neural timescales follows the spatial core-periphery organization. Data analyses including rest-task correlation and rest-task differences were complemented by kernel density estimation, machine learning, and simulation for a more precise differentiation of core and periphery regions on temporal grounds, i.e., their intrinsic neural timescales.

**Figure 1.**
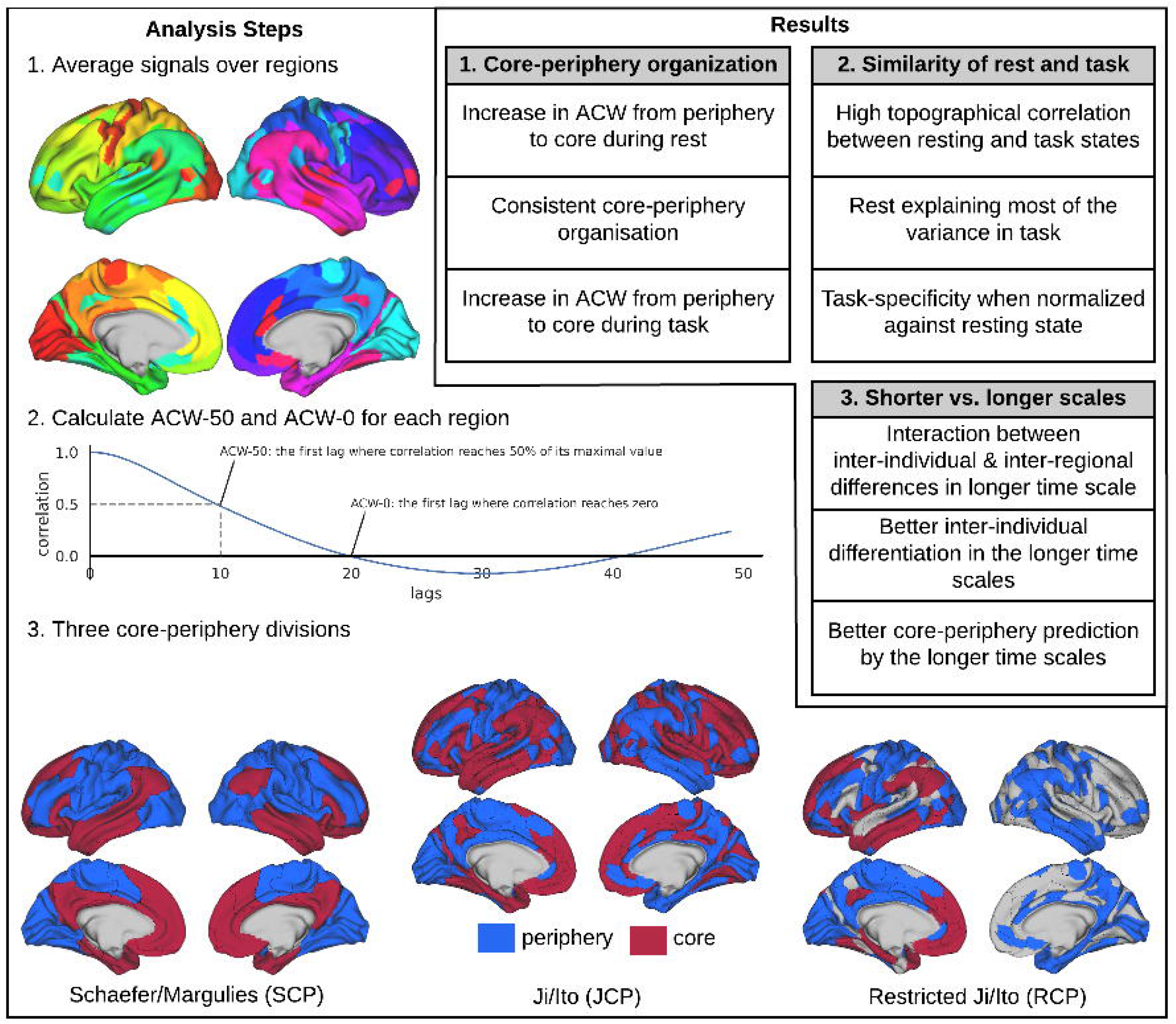
Schema of the paper. Represents the primary analysis conducted in this study. First, the time-series were averaged over cortical brain regions (1). Then, ACF was calculated for each region and from that ACW-50 and ACW-0 were extracted (2). Three different core-periphery divisions were defined based on the Schaefer and Ji templates (3). In Schaefer/Margulies (SCP) core-periphery division, cortical regions/networks are defined based on the Schaefer template ^40^ and then divided into core or periphery based on the first principal gradient introduced in ref. ^14^. The regions/networks in Ji/Ito (JCP) are from Ji template ^41^ and each region is labelled core or periphery based on unimodal/transmodal definition in ref. ^1^. The periphery definition in Restricted Ji/Ito (RCP) is similar to JCP, but only the regions in cingulo opercular, frontoparietal and default-mode networks are assigned to the core division, and the rest of the regions are ignored.

Intrinsic neural timescales can be measured by the autocorrelation window (ACW), which measures repeating patterns in a signal and enables testing for correlation in neural activity patterns at different points in time ^4^. The ACW has been determined at the cellular level ^4,5,34^ as well as systems-level (using fMRI ^1,3^ and EEG/MEG ^9,26,35^). Therefore, we here use ACW to measure the spatial CP organization in the faster frequency range of MEG. To account for different window sizes, we, in addition to the standard measurement of autocorrelation decline at 50% (ACW-50), also used a longer window size where ACW was defined as the length of time at the first instance where the Pearson correlation reaches zero (ACW-0) (See methods for details and differences between the two measures). Employing short (ACW-50) and long (ACW-0) window sizes enabled us to probe a more fine-grained temporal organization, that is, whether core and periphery regions can be differentiated by differences in the shorter and longer temporal windows within the faster frequency range of MEG (1.3-50Hz).

Our first specific aim focuses on the resting state as to investigate whether the temporal hierarchy of intrinsic neural timescales follows the spatial hierarchy of core-periphery organization. fMRI (0.01 to 0.1Hz) resting state studies of intrinsic neural timescales demonstrate short ACW in unimodal sensory regions and long ACW in transmodal regions ^1–3^ which suggests a CP organization. Extending these results to the faster frequency range (1.3-50Hz), we hypothesize that the topographical distribution of the resting state’s intrinsic neural timescales, i.e., ACW-50 and ACW-0, converges with the spatial CP organization – this would imply a truly temporo-spatial core-periphery hierarchy.

The second specific aim is to investigate the intrinsic neural timescales during different task states including their relation to resting state, i.e., rest-task correlation and rest-task differences. This is based on the findings, that rest and task data by themselves show strong hierarchical similarities in intrinsic neural timescales ^1,3,6,9,11^ as well as previous data showing that the core-periphery organization holds during task states ^17,36^. Together, these findings suggest carry-over of temporal and spatial hierarchies from rest to task states which we measure by rest-task correlation and rest-task differences. Taken together, we hypothesize that the convergence of temporal hierarchy with the spatial core-periphery organization also holds during task states as being carried over from rest to task; we, therefore, hypothesize a high correlation of the resting state’s spatial topography of the intrinsic neural timescales with the one during task states.

The third specific aim consists of probing whether core and periphery regions differ from each other by the length of the windows of their intrinsic neural timescales as measured by the different window sizes of ACW-50 (shorter window) and ACW-0 (longer window). This is based on the observation that, compared to the periphery, core regions/networks exhibit stronger slow frequencies ^37,38^ which leads to longer time windows ^9,39^ as measured with especially the ACW-0 (rather than the ACW-50). We, therefore, hypothesized that the ACW-0, probing longer time windows, allows differentiating core and periphery regions better than the shorter time window of ACW-50. To support the empirical data, we also use machine learning and simulation to verify this hypothesis that the longer time window of ACW-0 allows for more precise characterization of core region relative to periphery regions when compared to the shorter ACW-50.

## Results

In this paper, our main aim was to investigate whether the temporal hierarchy of intrinsic neural timescales of the faster frequency (1-50Hz) of MEG conform to core-periphery (CP) architecture and how they’ll be affected in the task state. Preprocessed MEG data of 89 subjects from the Human Connectome Project (HCP) were low-pass filtered at 50 Hz (thus the frequency range was 1.3-50Hz). One resting state (Rest, 6 minutes) and three task conditions of language processing (StoryM, 7 min), motor (Motort, 14 min), and working memory (Wrkmem, 10 min) were recorded from each subject. Sensor signals were projected onto source space by synthetic aperture magnetometry method and then parcellated using two well-known templates provided by Schaefer et al. ^40^ and Ji et al. ^41^. In the first parcellation (Schaefer) the surface-based data were parcellated into 200 regions (7 networks: Visual, Somatomotor, Dorsal Attention, Salience, Limbic, FPC and DMN) and 360 regions in the second (Ji) parcellation (12 networks: Visual1, Visual2, Auditory, Somatomotor, Dorsal Attention, Posterior Multimodal, Ventral Multimodal, Orbito Affective, Language, Cingulo Opercular, FPC and DMN).

Three different core-periphery divisions were defined in this work. 1) Schaefer/Margulies core-periphery (SCP): the regions from the Schaefer template were divided into core and periphery based on the principal gradient introduced in ref. ^14^ in which regions in the Limbic, FPC and DMN networks were put into the core category and the others in the periphery one. 2) Ji/Ito core-periphery (JCP): the regions from the Ji template were used for this CP definition. They were labelled core or periphery based on the unimodal/transmodal definition of ref. ^1^ in which regions in Visual1, Visual2, Auditory, and Somatomotor were assigned to the periphery, and the rest of the brain to the core. 3) Restricted Ji/Ito core-periphery (RCP) which was similar to JCP, but with a more restricted core definition. Only the regions of Cingulo Opercular, FPC and DMN networks were put into the core of RCP and the rest of the regions (not in either periphery or core) were ignored. The primary analysis steps are illustrated in Figure 1 and the findings are presented in the box of the same figure.

### ACW during resting state

In the first analysis, we calculated ACW values for each cortical region in the resting state data. For each region, two ACW scales were calculated, i.e. ACW-50 and ACW-0, as the first lags where autocorrelation function (ACF) reaches half of its maximum value, and zero, respectively. Next, the ACW values were averaged over subjects and were assigned to either core or periphery categories. All three CP divisions (Figures 2a for SCP, 2b for JCP and 2c for RCP) show an increase in both ACW scales from the periphery to the core. The difference between core and periphery was statistically tested using student’s t-test (with Cohen’s d for the effect size) which showed significant differences (*p* < 0.001) between the two in SCP (ACW-50: *t* = −5.25, *d* = −0.77, ACW−0: *t* = −5.42, *d* = −0.78), JCP (ACW−50: *t* = −5.72, *d* = −0.66, ACW−0: *t* = −11.67, *d* = −1.33) and RCP (ACW−50: *t* = −6.12, *d* = −0.75, ACW−0: *t* = −13.58, *d* = −1.63).

**Figure 2.**
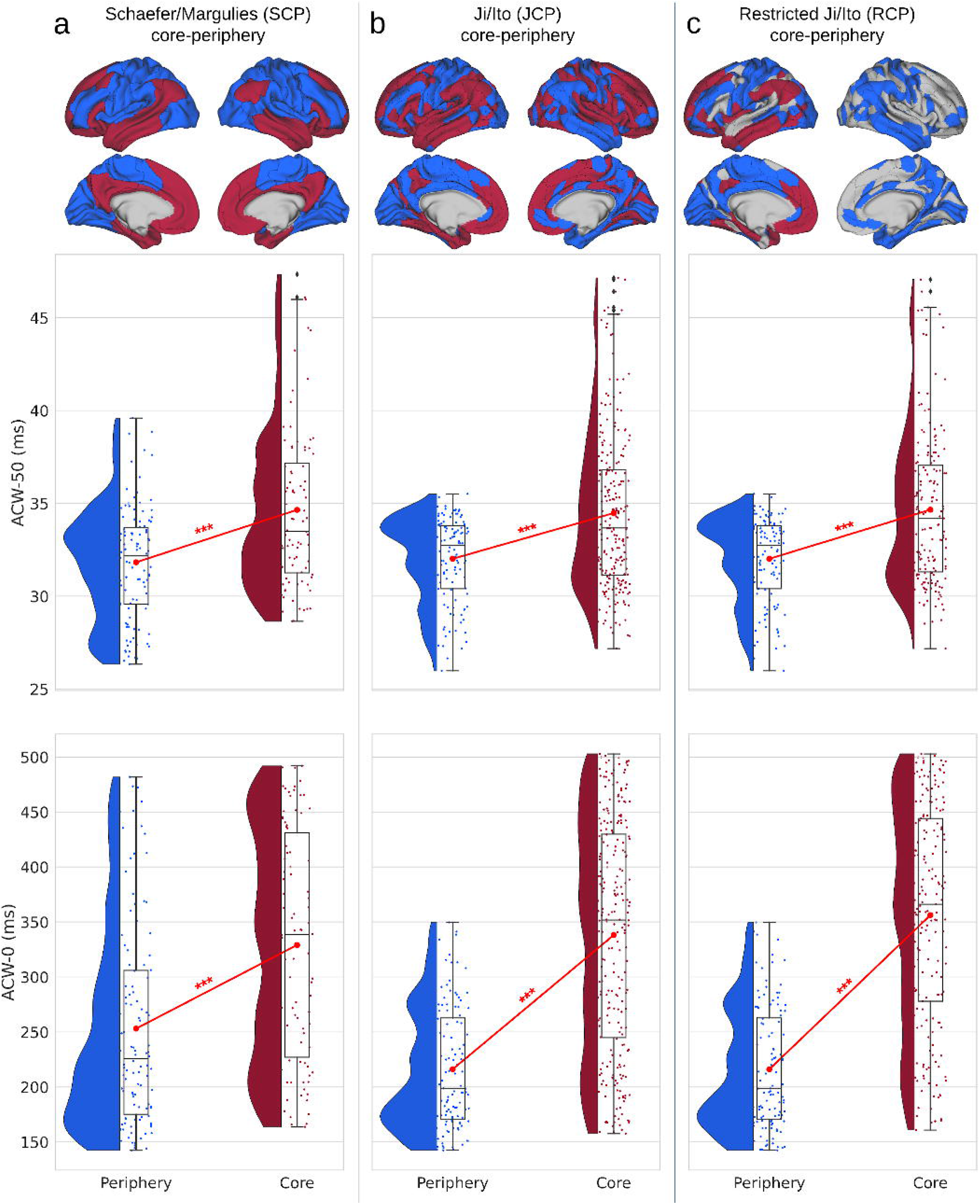
Autocorrelation window (ACW) during resting state for the three core-periphery (CP) divisions. Rainclouds represent regions. Values are presented in milliseconds. Brain plots show the different CP divisions of the regions. (a) Showing rainclouds for the ACW values of core and periphery using the Schaefer/Margulies division (SCP). The student’s t-test showed a significant difference in ACW-50 (*t* = −5.25, *d* = −0.77) and ACW-0 (*t* = −5.42, *d* = −0.78). (b) The same analysis as (a), but using the CP definition of the Ji/Ito (JCP). Student’s t-test was significant for the difference of both ACW-50 and ACW-0 between core and periphery (*t* = −5.72, *d* = −0.66 and *t* = −11.67, *d* = −1.33, respectively). (c) The same analysis using the restricted definition of core (RCP) showed a significant statistical difference between core and periphery for both ACW-50 (*t* = −6.12, *d* = −0.75) and ACW-0 (*t* = −13.58, *d* = −1.63). Stars represent the significance level (*** ≡ *α* = 0.001).

A recent article ^25^ argues that the peak frequency of the MEG signal in the range of 3-45 Hz follows the anterior-posterior axis (more details in the discussion). To further validate the core-periphery organization of ACW, we used linear regression to regress out the effect of this axis. An anterior-posterior value was calculated for each region based on the y coordinate of the region in the anatomical surface map (provided by the HCP group). Then, the anterior-posterior axis was used as the independent variable in the linear model of ACW. The residual of the model (residual ACW) is presented in the Supplementary Figure 1 for all three CP divisions. SCP (ACW-50: *t* = −7.80, *d* = −1.14, ACW-0: *t* = −7.23, *d* = −1.04), JCP (ACW-50: *t* = −10.43, *d* = −1.20, ACW-0: *t* = −11.65, *d* = −1.33), and RCP (ACW-50: *t* = −11.69, *d* = −1.43, ACW-0: *t* = −13.51, *d* = −1.62) show a significant (p < 0.001) difference between core and periphery of residual ACW, thus, validating our previous results.

The topography of ACW during resting state was further explored on a network level using both Schaefer (Supplementary Figure 2a) and Ji (Supplementary Figure 2b) templates. Plotting these values revealed different spatial patterns among the networks. One-way Analysis of Variance (ANOVA) suggests significant (*p* < 0.001) differences between networks in both ACW-50 (Schaefer: *F*(6,193) = 11.17,*η*^2^ = 0.25, Ji: *F*(11, 348) = 24.23, *η*^2^ = 0.43) and ACW-0 (Schaefer: *F*(6,193) = 21.42, *η*^2^ = 0.39, Ji: *F*(11, 348) = 215.64, *η*^2^ = 0.86).

### Organization of ACW along the principal gradient of core-periphery

Margulies et al. ^14^ suggest a principal gradient of functional connectivity for the core-periphery organization which captures the transition between different networks (Figure 3c from ref. ^14^). To further validate that the topography of ACW follows the core-periphery organization, we projected our results on to the same cutouts provided in Figure 3c of ref. ^14^ (Figure 3). The results suggest a direction of change similar to the principal gradient.

**Figure 3.**
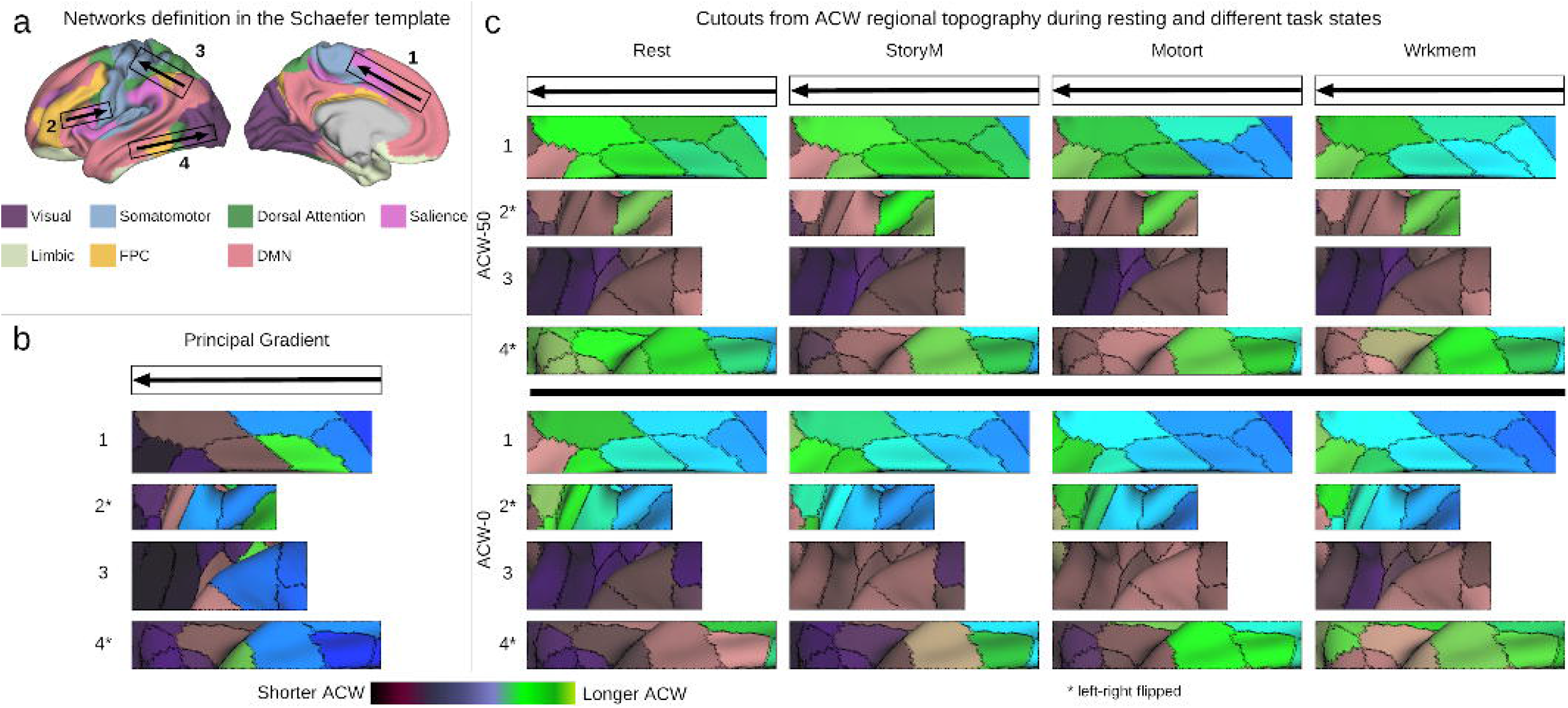
ACW cutouts for principal gradient of functional connectivity. The cutouts are similar to Figure 3c of ref. ^14^ They suggest that the direction of cross-regional changes in ACW is similar to the principal gradient of functional connectivity defined in ref ^14^ further validating the presence of the core-periphery organization in ACW. (a) Networks and the cutouts on the brain. (b) The regional principal gradient from the cutouts. (c) regional ACW values from the cutouts.

### ACW during task states

To investigate whether the CP organization of ACW is preserved during the task state, we calculated ACW-50 and ACW-0 for the three different task conditions of language processing (StoryM), motor (Motort) and working memory (Wrkmem). An increase in both ACW scales from the periphery to the core in all three task conditions was observed (Figure 4). To test that, we used three separate two-way Analysis of Variances (ANOVA) for the three CP organizations (SCP, JCP, and RCP). Both task (3 levels) and core-periphery (2 levels) were used as independent factors in the model. The model shows significant (*p* < 0.001) effect of core-periphery factor for SCP (ACW-50: *F*(1,594) = 57.69, *η*^2^ = 0.08, ACW-0: *F*(1,594) = 70.47, *η*^2^ = 0.09), JCP (ACW-50: *F*(1,1074) = 318.44, *η*^2^ = 0.22, ACW-0: *F*(1,1074) = 464.40, *η*^2^ =0.26), and RCP (ACW-50: *F*(1, 1074) = 311.75, *η*^2^ = 0.25, ACW-0: *F*(1,1074) = 549.90, *η*^2^ = 0.33). Complete result of the ANOVA test is presented in Supplementary Table 1. Further post-hoc analysis using Tukey’s HSD method reveals that regions in the periphery are significantly different from those in the core in all three CP divisions of all task conditions (Table 1). These results align with our resting state findings suggesting that the ACW in both 50 and 0 scales was higher in the core regions during all three tasks thus following a CP regime during task states.

**Figure 4.**
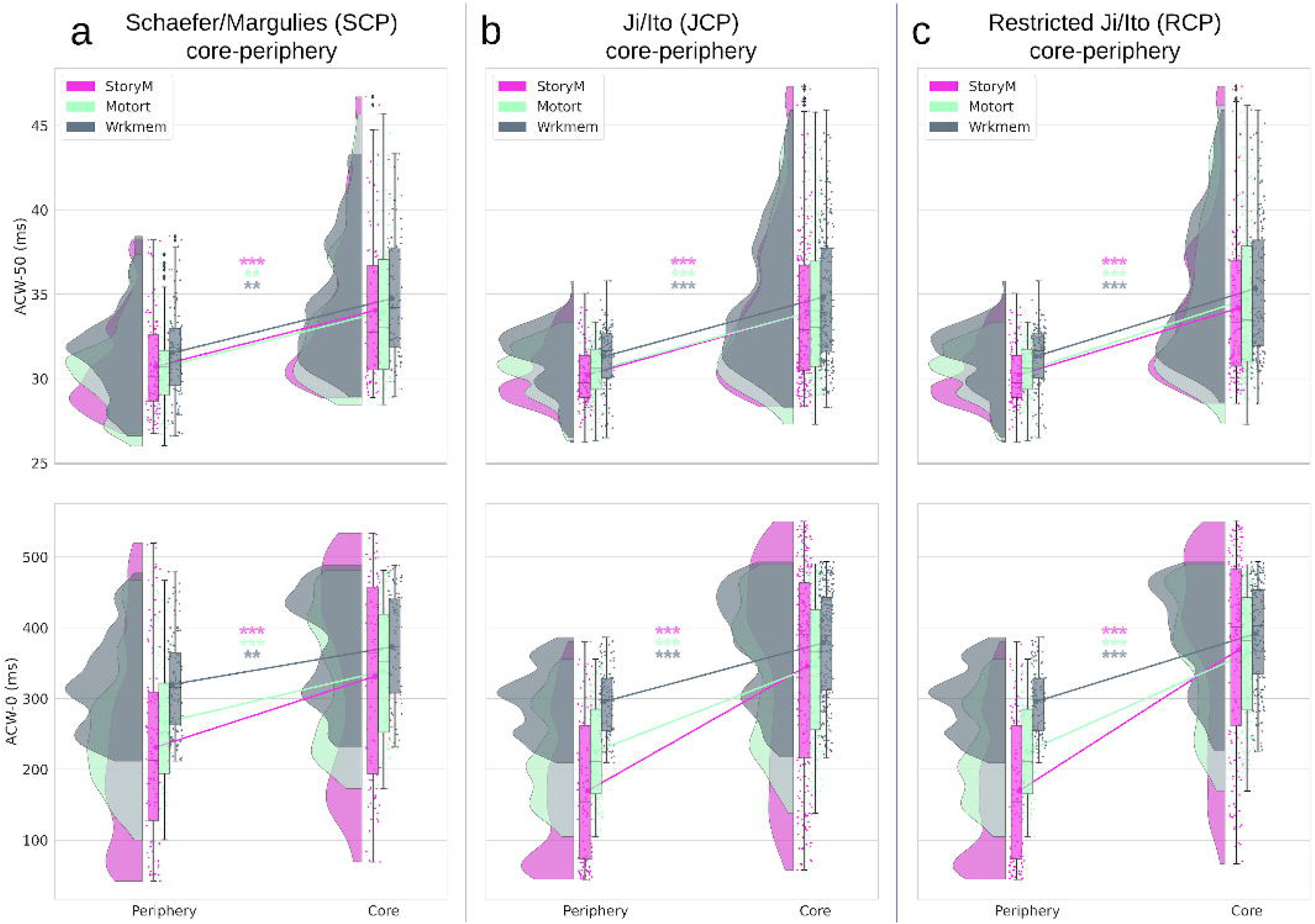
ACW during task conditions for different core-periphery divisions. Rainclouds represent regions divided into core and periphery. Values are presented in milliseconds. (a) Showing rainclouds for the ACW values of core and periphery using the Schaefer/Margulies division (SCP). Two-way analysis of variance (ANOVA) using task condition (3 levels: StoryM, Motort and Wrkmem) and CP (2 levels: core and periphery) as factors showed significant (p < 0.001) effect of CP factor on ACW-50 (*F*(1, 594) = 57.69, *η*^2^ = 0.08) and ACW-0 (*F*(1, 594) = 70.47, *η*^2^ = 0.09). Complete ANOVA result is presented in Supplementary Table 1. (b) Similar analysis as in (a) for the Ji/Ito core-periphery (JCP). Again, two-way ANOVA showed significant (*p* < 0.001) effect of CP factor on ACW-50 (*F*(1,1074) = 318.44, *η*^2^ = 0.22) and ACW-0 (*F*(1,1074) = 464.40, *η*^2^ = 0.26). (c) Similar to (a) and (b) but using the restricted core-periphery (RCP). Similar to previous results the effect of CP factor was significant (*p* < 0.001) for ACW-50 (*F*(1,1074) = 311.75, *η*^2^ = 0.25) and ACW-0 (*F*(1,1074) = 549.90, *η*^2^ = 0.33). Similar to the resting state results, an increase in both ACW scales along the different definitions of CP can be observed in all task conditions. Post hoc analysis using the Tukey HSD method revealed significant differences between periphery and core in all task conditions. The post-hoc results are presented in Table 1. Stars represent the significance level (*** ≡ *α* = 0.001,** ≡ *α* = 0.01).

**Table 1.**
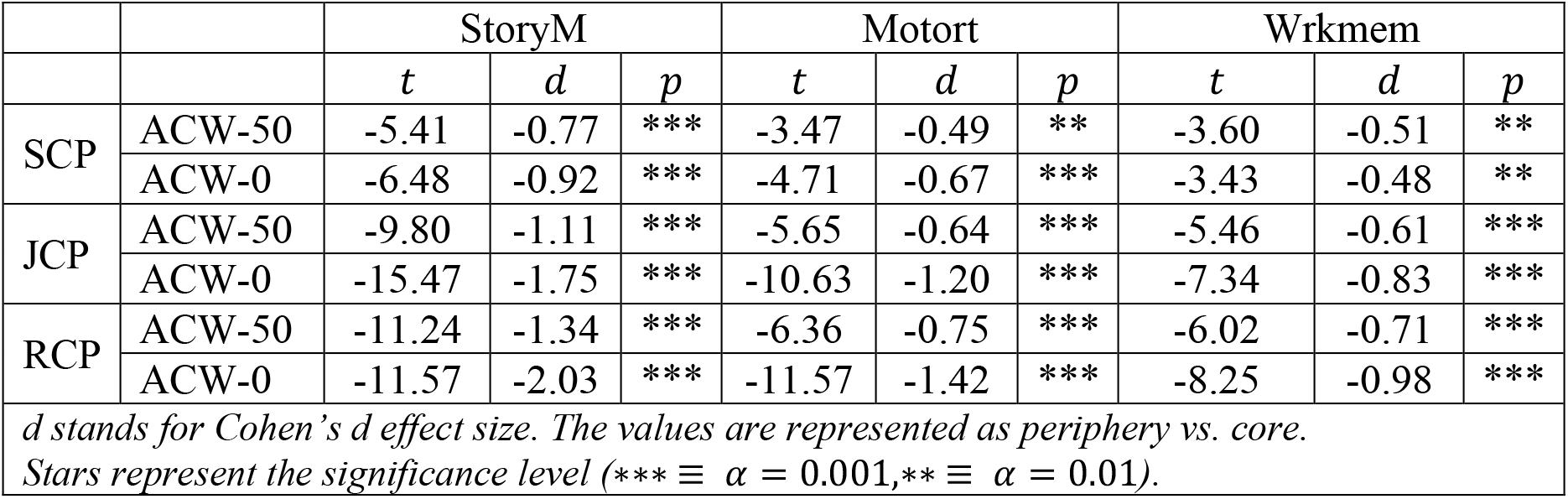
Post-hoc results for periphery vs. core in different task conditions. Tukey HSD method was used to determine the between factor significance of ANOVA on both ACW-50 and ACW-0 along different CP divisions.

|  |  | StoryM |  |  | Motort |  |  | Wrkmem |  |  |
| --- | --- | --- | --- | --- | --- | --- | --- | --- | --- | --- |
|  |  | *t* | *d* | *p* | *t* | *d* | *p* | *t* | *d* | *p* |
| SCP | ACW-50 | -5.41 | -0.77 | *** | -3.47 | -0.49 | ** | -3.60 | -0.51 | ** |
|  | ACW-0 | -6.48 | -0.92 | *** | -4.71 | -0.67 | *** | -3.43 | -0.48 | ** |
| JCP | ACW-50 | -9.80 | -1.11 | *** | -5.65 | -0.64 | *** | -5.46 | -0.61 | *** |
|  | ACW-0 | -15.47 | -1.75 | *** | -10.63 | -1.20 | *** | -7.34 | -0.83 | *** |
| RCP | ACW-50 | -11.24 | -1.34 | *** | -6.36 | -0.75 | *** | -6.02 | -0.71 | *** |
|  | ACW-0 | -11.57 | -2.03 | *** | -11.57 | -1.42 | *** | -8.25 | -0.98 | *** |
| *d* stands for Cohen's *d* effect size. The values are represented as periphery vs. core.<br>Stars represent the significance level (*** $\equiv \alpha = 0.001$, ** $\equiv \alpha = 0.01$). | | | | | | | | | | |

To extend the resting state results and further validate the CP organization of ACW scales during task conditions, we controlled for the anterior-posterior gradient by regressing out the y coordinate of each region and repeating the aforementioned analysis on the residual ACW values. The y coordinate of all regions was used as the independent variable in a linear regression model (one model per each scale) to estimate the ACW values (ACW values used as dependent variables). Then, the residual of the model was determined as the residual ACW and plotted along the three definitions of CP, i.e. SCP, JCP, and RCP (Supplementary Figure 3). Statistical analysis using two-way ANOVA (similar to previous results, complete ANOVA in Supplementary Table 2) showed a significant (*p* < 0.001) effect of the core-periphery factor on residual ACW. SCP: ACW-50, *F*(1,594) = 152.67, *η*^2^ = 0.20 ; ACW-0: *F*(1,594) = 144.40, *η*^2^ = 0.19). JCP: ACW-50, *F*(1,1074) = 514.64, *η*^2^ = 0.32; ACW-0, *F*(1,1074) = 518.48, *η*^2^ = 0.32. RCP: ACW-50, *F*(1,1074) = 565.43, *η*^2^ = 0.38; ACW-0: *F*(1,1074) = 607.97, *η*^2^ = 0.40. Moreover, post-hoc analysis using Tukey’s HSD method revealed that the periphery has significantly lower ACW values compared to the core in the task condition within all three different definitions of CP (Supplementary Table 3). These results validate our original results suggesting that ACW during task, similar to rest, follows a core-periphery organization.

Task state ACW was also investigated along the different networks defined by both Schaefer and Ji templates. Both ACW scales were first averaged over subjects and then plotted for different networks (Supplementary Figure 4). The effect of networks on the ACW values was tested using ANOVA. First, ACW was modelled with both task (3 levels) and network (6 levels in Schaefer and 12 levels in Ji templates) as independent factors to test if the effect of network is overall significant. Then, a one-way ANOVA with only the network as the independent factor was calculated for each task to determine the effect of network in each task. The results are presented in Supplementary Table 4 showing a significant (*p* < 0.001) effect of network in both templates on ACW-50 and ACW-0 during all three task conditions.

### The relationship between resting and task states in ACW

#### The similarity between resting and task states

In this step, we investigated how the resting state’s intrinsic neural timescales including their CP-gradient are carried over to task states; for that, we conducted various analyses on rest-task similarities and differences. The rest-task similarity analysis was performed in three steps, spatial correlation, linear regression and regional correlation.

The spatial correlation was calculated to see how similar the spatial distribution of ACW values are between resting and different task states (Figures 5a for ACW-50 and 5b for ACW-0). It was conducted by first averaging each condition (either resting or task) across subjects and then correlating resting state’s averaged brain map with each task state’s brain map across regions. Pearson correlation showed that all tasks ACW’s spatial distributions are highly (n = 360, *p* < 0.001) correlated with the resting state in both ACW scales, i.e. 50 and 0, (values are presented in Figures 5a and 5b) suggesting similar spatial topography between resting and task states.

**Figure 5.**
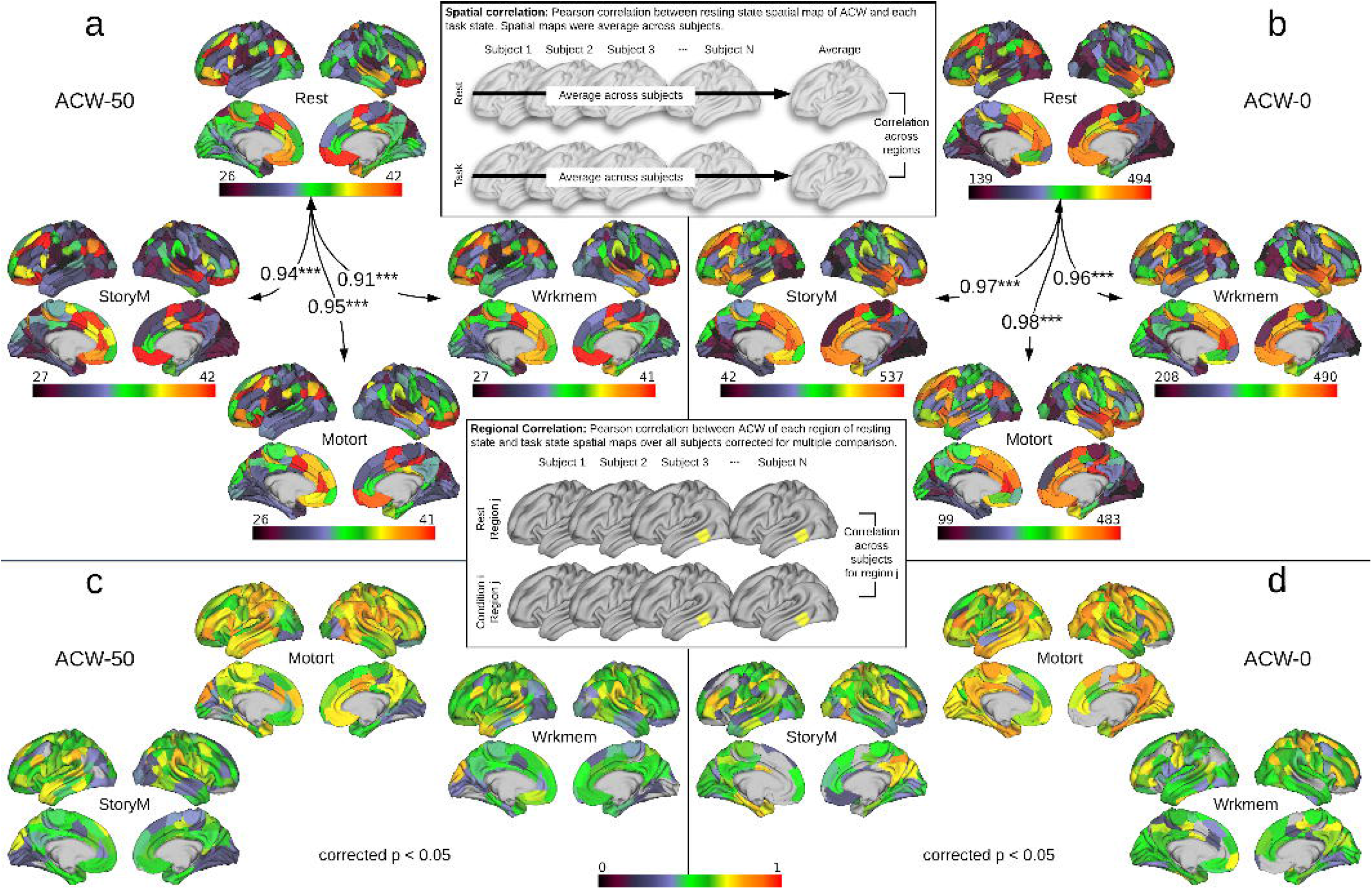
The spatial similarity between resting and task states. The similarity was calculated by applying rest-task spatial correlation (a, b) and intra-regional rest-task correlation (c, d). The boxes with the gray brains show how spatial and regional correlation was calculated. The yellow region in the box of Regional Correlation is a sample region chosen only for illustration purpose. (a) and (b) show spatial correlation (numbers on the arrows) between different conditions alongside their spatial distribution over brain regions for ACW-50 and ACW-0, respectively. Each value on the arrows is a Pearson correlation coefficient between a pair of conditions across the brain regions (*n* = 360). All correlation values suggest that different conditions are highly correlated with each other (*** ≡ *α* = 0.001). (c) and (d) show the regional correlation between resting and each task state in ACW-50 and ACW-0, respectively. Pearson correlation coefficients were calculated for each region over different subjects, then corrected for multiple comparisons using the FDR correction method at *α* = 0.05. Only the regions that survived the correction are illustrated.

To extend our spatial correlation results, we also conducted linear regression. ACW during each task state was modelled as a function of the resting state in a linear model (Supplementary Figure 5) after averaging all conditions (either resting or task) across subjects (thus, one brain map for each condition). This analysis revealed a significant (n = 360, *p* < 0.001) linear relationship between Rest and all three task conditions in both Schaefer and Ji parcellation templates (Æ^2^ values are presented in Supplementary Figure 5) suggesting that rest ACW can explain more than 85% of the variance in task ACW.

Next, the rest-task similarity was further explored using regional correlation. The results were analogous for both ACW-50 (Figure 5c) and ACW-0 (Figure 5c). In both cases, first, a Pearson correlation coefficient was calculated for each region over different subjects (*n* = 89) between resting and each task state (e.g. Rest vs. StoryM). Then, p-values were corrected for multiple comparisons using the False Discovery Rate (FDR) method at *α* = 0.05 while non-significant coefficients were ignored. The results show a high correlation of resting and task states across subjects suggesting that the similarity between resting and task states is consistent across subjects. Together, the findings show strong rest-task similarity suggesting that the convergence of temporal and spatial hierarchies during the resting state is carried over to task states.

#### Degree of change from resting to task state

This analysis aimed to investigate the degree of task-related changes relative to rest and whether that change also conforms to a CP regime. For that, we calculated the percentage of change from resting to task state. It was calculated per region and per subject in all three task conditions and was normalized against their respective resting state values, i.e., the degree of change from rest to task (see methods for details), in all three CP divisions (SCP, RCP and JCP, Figure 6). The percentage of change was then tested for statistical significance using two-way ANOVA similar to previous resting and task state analyzes. Task condition (3 levels) and core-periphery (2 levels) were used as independent factors to investigate their effects on the percentage change of ACW scales from rest to the task. The statistical test (Supplementary Table 5) suggests a significant (*p* < 0.001) effect of core-periphery factor on the change in ACW in SCP (ACW-50: *F*(1, 594) = 10.63, *η*^2^ = 0.01, ACW-0: *F*(1, 594) = 12.17, *η*^2^ = 0.01), JCP (ACW-50: *F*(1,1074) = 99.61, *η*^2^ = 0.07, ACW-0: *F*(1,1074) = 26.74, *η*^2^ = 0.01), and RCP (ACW-50: *F*(1,1074) = 97.35, *η*^2^ =0.08, ACW-0: *F*(1,1074) = 34.92, *η*^2^ = 0.02). The difference between periphery and core was also investigated using Tukey’s HSD method in a post-hoc analysis (Table 2). We can see a prominent change in ACW from rest to task. However, the direction of change is not similar among the three conditions (e.g. StoryM and Motort showing opposite directions of change from rest to task in ACW-0). These findings suggest task-specific effects in especially ACW-0 but also in ACW-50. Notably, these task-specific findings were not observed in the task conditions alone, i.e., independent of rest (Figure 4); they become only apparent when one subtracts task from rest, i.e., rest-task difference, which subtracts, i.e., eliminates, those features shared by rest and task, e.g., core-periphery organization.

**Figure 6.**
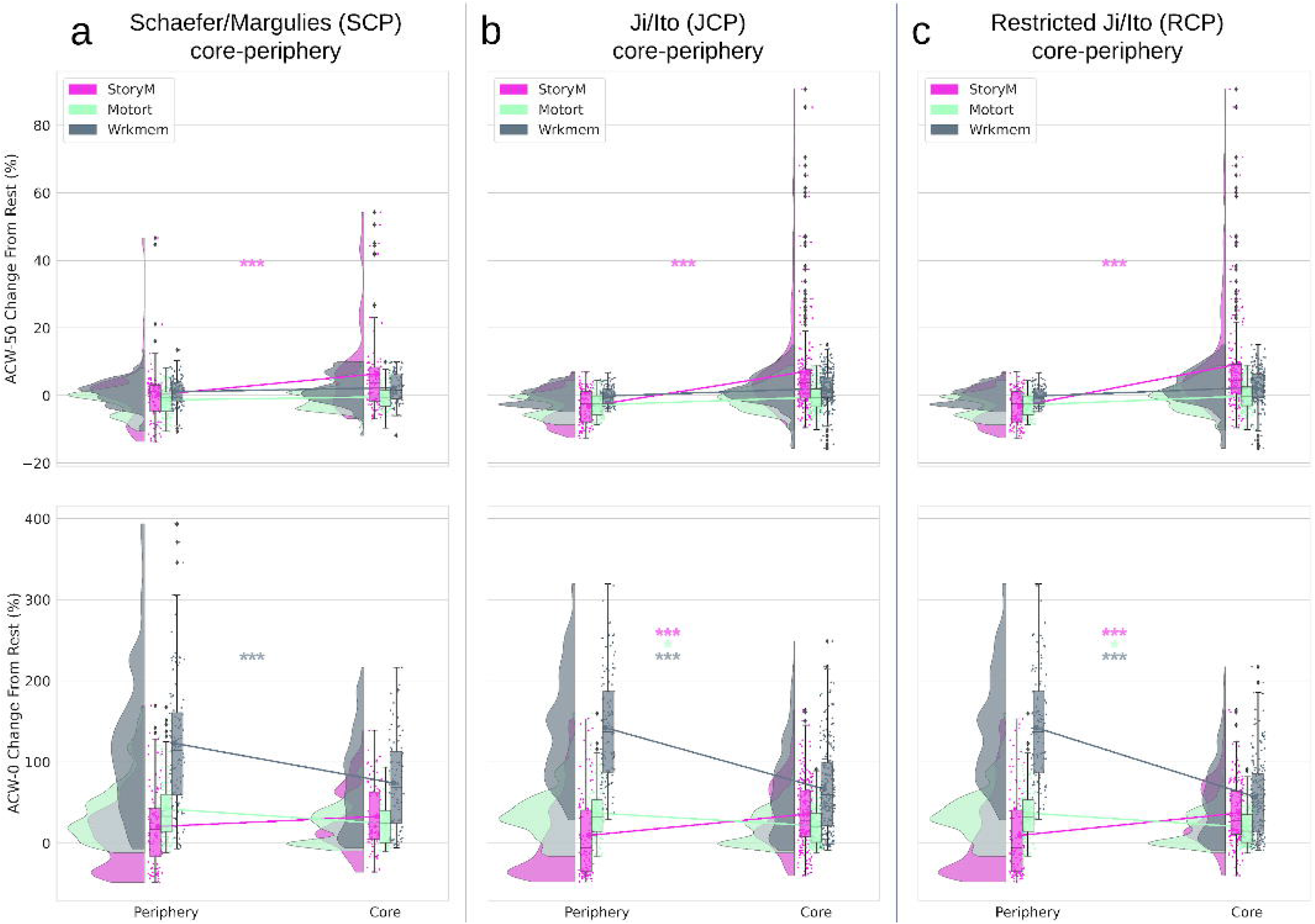
Change in ACW from rest to task states in different CP divisions by calculating rest-task differences. Rainclouds represent regions divided into core and periphery. Values are presented in percentages. Stars represent the significance level (*** ≡ *α* = 0.001,* ≡ *α* = 0.05). (a) Showing rainclouds for the ACW change values of core and periphery using the Schaefer/Margulies division (SCP). A statistical test (Supplementary Table 5) was performed with two-way ANOVA using task condition (3 levels: StoryM, Motort and Wrkmem) and CP (2 levels: core and periphery) as independent factors. It showed significant (*p* < 0.001) effect of CP factor on ACW-50 (*F*(1,594) = 10.63, *η*^2^ = 0.01) and ACW-0 (*F*(1,594) = 12.17, *η*^2^ = 0.01). (b) Ji/Ito template (JCP): again, two-way ANOVA showed significant (*p* < 0.001) effect of CP factor on ACW-50 (*F*(1,1074) = 99.61, *η*^2^ = 0.07) and ACW-0 (*F*(1,1074) = 26.74, *η*^2^ = 0.01). (c) Restricted Ji/Ito template (RCP): significant (*p* < 0.001) effect of CP for ACW-50 (*F*(1,1074) = 97.35, *η*^2^ = 0.08) and ACW-0 (*F*(1,1074) = 34.92, *η*^2^ = 0.02). Post-hoc analysis was performed using Tukey’s HSD method to determine whether the difference between core and periphery is significant. The results are provided in Table 2.

**Table 2.**
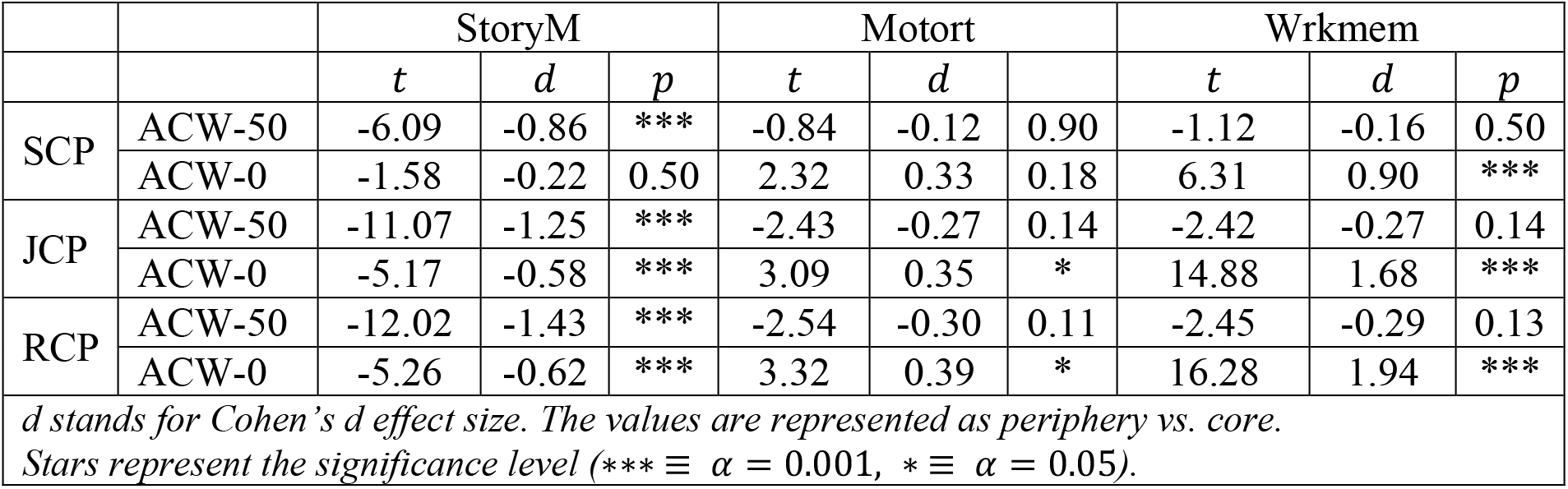
Post-hoc results for periphery vs. core in the change from resting to task state. Tukey’s HSD method was used to determine the between factor significance of ANOVA on both ACW-50 and ACW-0 along different CP divisions.

|  |  | StoryM |  |  | Motort |  |  | Wrkmem |  |  |
| --- | --- | --- | --- | --- | --- | --- | --- | --- | --- | --- |
|  |  | *t* | *d* | *p* | *t* | *d* |  | *t* | *d* | *p* |
| SCP | ACW-50 | -6.09 | -0.86 | *** | -0.84 | -0.12 | 0.90 | -1.12 | -0.16 | 0.50 |
|  | ACW-0 | -1.58 | -0.22 | 0.50 | 2.32 | 0.33 | 0.18 | 6.31 | 0.90 | *** |
| JCP | ACW-50 | -11.07 | -1.25 | *** | -2.43 | -0.27 | 0.14 | -2.42 | -0.27 | 0.14 |
|  | ACW-0 | -5.17 | -0.58 | *** | 3.09 | 0.35 | * | 14.88 | 1.68 | *** |
| RCP | ACW-50 | -12.02 | -1.43 | *** | -2.54 | -0.30 | 0.11 | -2.45 | -0.29 | 0.13 |
|  | ACW-0 | -5.26 | -0.62 | *** | 3.32 | 0.39 | * | 16.28 | 1.94 | *** |
| *d* stands for Cohen's *d* effect size. The values are represented as periphery vs. core.<br>Stars represent the significance level (*** $\equiv \alpha = 0.001$, * $\equiv \alpha = 0.05$). | | | | | | | | | | |

Taken together, our results, on one hand, suggest a consistent similarity in the organization of ACW between resting and different task states – this is documented by their high similarity in all three statistical rest-task analyses. On the other hand, when the effect of resting state is removed, i.e., eliminated, as when calculating the rest-task differences, each task shows distinct features in its temporal organization. These results suggest that task-specific effects occur within the framework of the more fundamental core-periphery organization as that is preserved during both rest and task states.

### The relationship between different timescales and their predictive power

So far, we have investigated the temporal organization of neural activity using two different scales of ACW, i.e. 50 and 0 which showed more or less similar results for both. The last part of our analysis involves investigating the relationship between ACW-50 and ACW-0 and showing the differential effects of them in determining core-periphery organization in a more fine-grained way. This was done using three methods: kernel density estimation, simulation and machine learning.

#### Kernel density estimation

First, the density estimation of the ACW data was performed to observe the general density function (DF) over the whole data. We used our Ji-parcellated data (360 regions per subject) and removed all the labels (e.g. task, subject, region) from the ACW values; then the range of each ACW scale (i.e. 50 or 0) was scaled to be between 0 and 1. Based on the histograms of the data, we decided to use the Gaussian kernel with the expectation-maximization algorithm to estimate the two DFs (for ACW-50 and ACW-0). Figure 7a left shows the estimated DFs. It can be observed that the range of ACW-0 values is much wider than ACW-50; this suggests a higher variety or range of ACW-0 values in the probability space which may allow for a more finegrained differentiation between different labels (i.e., condition, subject, or region).

**Figure 7.**
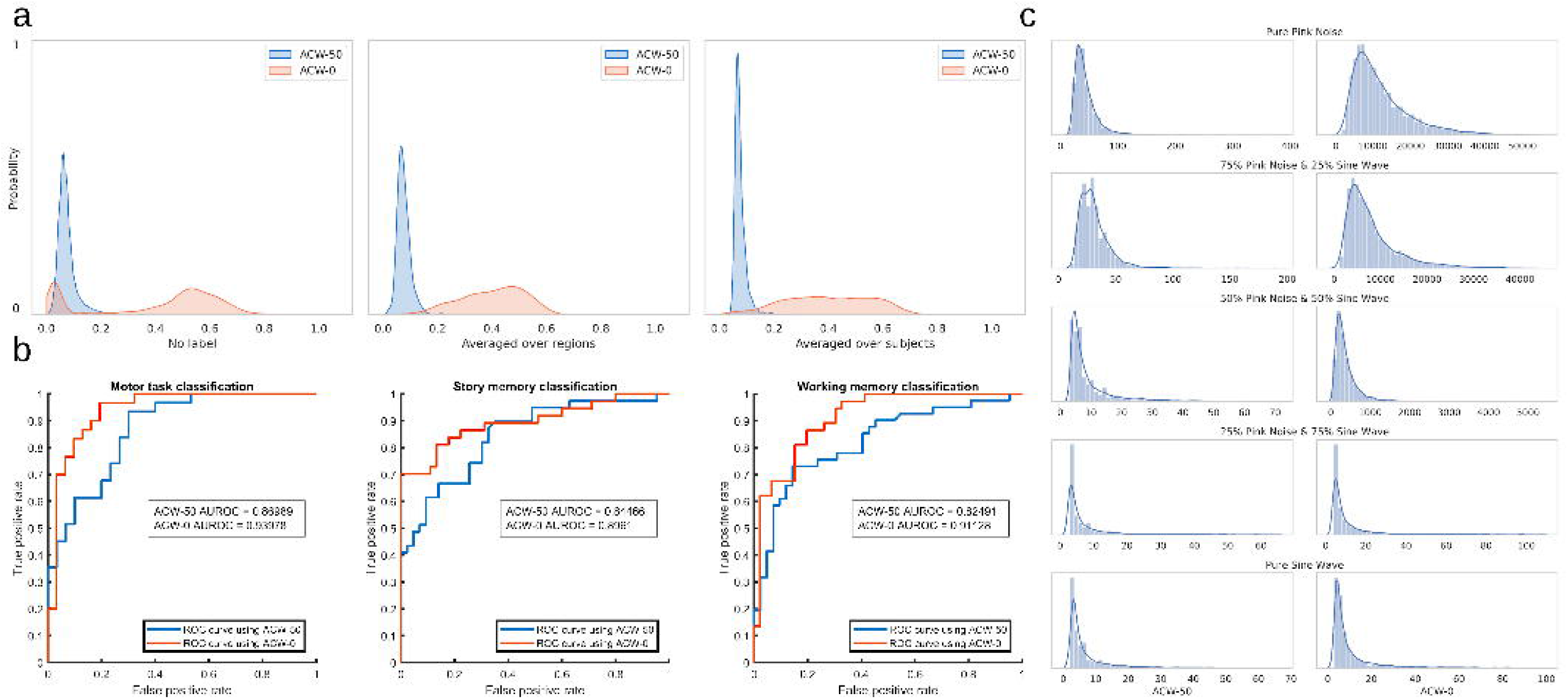
Relationship between shorter (ACW-50) and longer (ACW-0) time window measures. (a) The probability density of the data removing all labels (condition, subject or region), averaged over the region and averaged over the subjects. The Ji-parcellated data (360 regions per subject) was used for this analysis. The range of the data is scaled between 0 and 1 separately for ACW-50 and ACW-0. The density estimation with no label suggests a single narrow Gaussian distribution for ACW-50 and two separate Gaussians for ACW-0. When averaged over either regions or subjects, the two Gaussians of ACW-0 disappears. Thus, the double distribution of ACW-0 is only presented when the interaction of inter-individual differences with regional differences is taken into consideration. Moreover, ACW-0 is expanded over a wider range of values compared to ACW-50 suggesting better and more fine-grained differentiation capacity in the probability space. (b) Receiver operating characteristic (ROC) curves for discriminating subjects in different task conditions (StoryM, Motort, and Wrkmem). They suggest that ACW-0 has better discriminative power among subjects than ACW-50. (c) The density estimation of ACW values in simulated signals.

Importantly, the data suggest that ACW-0 follows two separate Gaussian distributions. To investigate whether these two distributions are related to subjects or/and regions, we repeated the same analysis once averaging the data over the regions to remove the effect of the inter-regional differences (Figure 7a middle) and once averaging over the subjects to remove the effect of the inter-individual difference (Figure 7a right). This revealed that the two separate DFs of ACW-0 only appear when we take the interaction of inter-individual and inter-regional differences (subject X region interaction) into consideration (the DFs without any label, i.e. Figure 7a left). That observation was consistent in all rest and task conditions suggesting that some subjects’ regional ACW follows the first DF and some subjects follow the second one. Together, these findings suggest considerable inter-subject variability in specifically the longer timescale, i.e., ACW-0, whereas inter-subject variability was rather minimal in the shorter timescale, i.e., ACW-50.

The impact of inter-individual differences in ACW-50 and ACW-0 was further validated using a binary classification problem. First, ACW values were averaged over regions (one ACW value per subject) and divided into two groups of “low variability” and “high variability”. The median of ACW values during resting state was used as the dividing threshold. Thus, the subjects were divided and labeled (actual label) as “low variability” or “high variability” considering only the activity during resting state. Second, a classification analysis was carried out on the activity during different tasks using a logistic regression analysis. For each task, the predicted labels (output of the logistic regression) were compared to actual ones (created from the resting state data in the first step) across all subjects. The area under the Receiver Operating Characteristic (ROC) curve was computed based on the predicted and actual labels (Figure 7b). An ROC curve is a graphical plot that illustrates the diagnostic ability of a binary classifier system as its discrimination threshold is varied. The area under the ROC curve (AUROC) tells how much the model is capable of distinguishing between the two classes. The higher the area under the curve, the better the model is at correctly predicting the true label. Results show that all the obtained AUROCs from ACW-0 are higher than those from ACW-50 for the three tasks. This implies that the ACW-0 contains more information than ACW-50 or, in other words, the predictive capability of ACW-0 to discriminate between subjects of high and low variability during the tasks are higher than the ACW-50.

#### Simulation

In the second analysis, we were interested to see if the double Gaussian distribution of ACW-0 is due to random noise or is indeed related to the brain signal. For that, we simulated random oscillatory and noise signals in distinct frequencies as distinguished by their different kinds of noises. Pink noise and sine waves were used as the basis of our pseudo-aleatory signals. They were simulated in the same frequency range and sampling rate as our original data (1.3-50 Hz) with uniformly distributed random weights. 20000 signals categorized into four distinct categories (5000 each) were used: pure pink noise, pure sine wave, a combination of sine and pink noise, and a combination of sine, pink and white noise. Random uniform coefficients were used to linearly combine signals in the third and fourth categories (see Supplementary Figure 6 for a sample signal in each category). Pink noise was utilized to simulate scale-freeness ^42^, sine wave for oscillation and white noise for pure randomness. Both ACW scales were calculated for the signals and plotted against each other in Supplementary Figure 6. Kernel density of each category was also estimated (Figure 7c); however, none of the simulated signals yielded the specific distributions like the ones previously observed in the brain signal suggesting that the two Gaussians observed in ACW-0 is not random, validating our previous results.

#### Machine learning

As the third and final analysis, we used machine learning to determine how the two ACW scales differ in predicting whether a region belongs to the core or periphery. To simplify the question without losing generality, we designed a two-class classification problem using a logistic classifier to determine if a signal is from a core (class 1) or periphery (class 2) region. Table 3 summarizes the results which show that not only the two scales have different performances, but also ACW-0 is a better predictor than ACW-50. The results were also replicated with a support vector machine (SVM) classifier using the radial basis function (RBF) kernel (Table 3).

**Table 3.** The results for the 2-class classification problem. The values are averaged over 20 folds of the K-fold cross-validation algorithm using Logistic regression and support vector machine (SVM) classifiers. The results are presented if each sample has two features (both ACW-50 and ACW-0), or only one feature (either ACW-50 or ACW-0). They suggest better classification power for ACW-0 compared to ACW-50.

| Algorithm | Model # | Features | Mean Accuracy (%) | Mean Precision (%) | Mean Recall (%) | Mean AUC of ROC |
| --- | --- | --- | --- | --- | --- | --- |
| Logistic Regression | 1 | ACW-50 & ACW-0 | 69.14 | 63.33 | 46.61 | 74.12 |
|  | 2 | Only ACW-50 | 60.55 | 40.61 | 6.02 | 61.10 |
|  | 3 | Only ACW-0 | 69.08 | 63.37 | 46.12 | 74.45 |
| SVM | 1 | ACW-50 & ACW-0 | 69.15 | 63.14 | 47.20 | 73.19 |
|  | 2 | Only ACW-50 | 61.61 | 41.21 | 7.01 | 48.72 |
|  | 3 | Only ACW-0 | 69.13 | 63.12 | 47.07 | 70.68 |

Next, we used a univariate feature selection method to determine which feature (ACW-50 or ACW-0) is better if we were required to choose one of the two. This method works by selecting the best features based on a univariate statistical test. We used mutual information as our scoring function ^43^ which is a data-driven method capable of capturing any kind of statistical dependency between two variables. It is equal to zero if and only if the class label is independent of the ACW scale (which means the scale is not a good feature) and higher values mean higher dependency between the two. It suggested that ACW-0 (score = 0.5) was a better feature compared to ACW-50 (score = 0.1) when predicting a region’s core or periphery association.

## Discussion

We demonstrate that the temporal hierarchy of intrinsic neural timescale measured by shorter and longer ACW, i.e., ACW-50 and ACW-0 follows the spatial topography namely the core-periphery hierarchy. Networks in the core exhibit longer intrinsic timescale, i.e., ACW-50 and ACW-0, than those at the periphery – this holds across both rest and task states as supported by strong rest-task correlation. Comparing rest and task reveals task-specific changes in particular networks only when the core-periphery organization is eliminated by subtraction, i.e., rest-task differences. Finally, we demonstrate that the longer timescale measure, i.e., ACW-0, exhibits higher accuracy in differentially classifying core vs periphery regions when compared to the shorter ACW-50; this suggests a stronger impact of slower frequencies in the core than in periphery regions resulting in their better and more precise differentiation. Together, we demonstrate that the temporal hierarchy of (shorter and longer) intrinsic neural timescales converges with the spatial topography of the core-periphery hierarchy of the human cortex with both providing an intrinsically temporo-spatial hierarchy during rest and task states.

### Core-periphery organization – converging spatial and temporal hierarchies

The spatial topography of the core-periphery hierarchy of the human cortex has been observed in resting state ^14–16,19^. We extend these findings beyond the spatial domain by showing a corresponding temporal hierarchy of intrinsic neural timescale. Animal ^4^ and modelling ^12,13,44^ studies observed different intrinsic neural timescales in lower- and higher-order regions/networks of the brain. This leaves open whether the human brain exhibits an analogous temporal architecture. Extending the animal and modelling findings to the human cortex, we, in rest states, observed shorter durations in ACW-50 and especially ACW-0 in periphery regions/networks like sensory and motor networks. That was complemented by longer ACW (especially in ACW-0) durations in the core networks like DMN, cingulum operculum, and FPN.

Together, these findings strongly suggest an intrinsic, i.e., network-specific, temporal architecture with different intrinsic neural timescales following the spatial topography, that is, the core-periphery hierarchy. One may consequently want to speak of an integrated temporospatial core-periphery hierarchy. Future investigation is needed to establish a more intimate link between temporal and spatial dimensions. For instance, one may raise the question of whether the functional connectivity among the regions of the core, i.e., core-core connectivity, is directly related to the longer intrinsic timescale in these regions as it is suggested by the recent findings of close relation between inter-regional functional connectivity and intra-regional ACW ^1,3^. Yet another example could be to link the core-periphery hierarchy to other temporal dynamic measures like the peak frequency, which recently has been shown to exhibit anterior-posterior hierarchy^25^. This raises the question, whether the anterior-posterior hierarchy is embedded in the more comprehensive core-periphery hierarchy which, by itself, was not affected when regressing the anterior-posterior gradient (See results).

An analogous temporo-spatial core-periphery hierarchy was observed during task states. Periphery networks showed shorter ACW while core networks exhibited longer ACW during all three task states. These findings suggest that the temporo-spatial core-periphery hierarchy is carried over from rest to task state holding across different tasks. One may consequently assume that the temporospatial core-periphery hierarchy may be an intrinsic feature that remains independent of the respective context, i.e., rest or task. That converges well with the observation that spatial core-periphery hierarchy has also been observed during task states ^17,36^ which may then be interpreted as simple carry-over from the rest. This remains to be explored in future studies.

Finally, findings on intrinsic neural timescale show higher-order networks like the DMN display the longest temporal receptive windows (based on functional connectivity) during equally complex tasks, i.e., story or movie ^6–9,11,45^. We replicated these findings in terms of ACW during task states and extend it to the rest where an analogous temporal hierarchy was observed. Moreover, when comparing rest and task ACW, i.e., rest-task differences, we observed task-specific effects that no longer followed the core-periphery hierarchy which we assume to be related to the cancelling out of the core-periphery hierarchy commonly shared by rest and task states. That, in turn, opened the door for observing task-specific ACW changes in particular networks. We demonstrate that the probability density of ACW shows a propensity towards longer ACW-0 in specifically the core networks while it remains shorter in periphery networks. This suggests some network-specific effects of longer (ACW-0) and also shorter (ACW-50) intrinsic neural timescale. Specifically, ACW-50 may be ideal to measure differences in shorter durations of intrinsic timescales in the periphery which are lost in the longer time window of ACW-0. Conversely, ACW-0 measuring longer time interval may be better suited to differentiate between regions with longer intrinsic timescales in specifically the core as it is suggested by our probability density function analysis.

### Rest shapes task states – intrinsic neural timescales enable temporal integration

Given that a core-periphery organization of intrinsic neural timescales in both rest and task states was observed, we tested for their relationship. We observed a high degree of correlation in the spatial differences for rest ACW-50/ACW-0 with their corresponding values in the three task states as well as among the three task states. That was complemented by high degrees in the correlation of the levels of ACW rest with those during the three task states. Together, these findings strongly support the assumption of an intrinsic temporal architecture in the brain that is preserved across rest and task states thus shaping both – the core-periphery organization is carried over from rest to task states independent of task-specific changes.

Our data suggest an intimate relationship between the intrinsic neural timescales during rest and task. We assume that such intimate relationship consists of temporal integration. The ACW and, more generally, the intrinsic neural timescales allow for temporal integration of stimuli through their temporal summing and pooling ^39^. The need for temporal integration is especially relevant in complex stimuli or tasks like the movie: complex sequences of stimuli need to be integrated here to apprehend their meaning. This requires a prolonged autocorrelation window in especially higher-order networks as it is suggested by our rest-task findings. Moreover, complex stimuli like music or movies involve multiple time scales including short and long – a corresponding hierarchy of timescales on the neural side may consequently be best for processing complex inputs ^11^.

In addition to the temporal integration of the various stimuli within the task itself, the external task-related stimuli need to be temporally integrated also with the spontaneous activity’s ongoing internal stimuli as related to internal cognition like mind-wandering ^46^ and bodily inputs ^47,48^. Such temporal integration may be mediated by the changes in the ACW during the transition from rest to task ^47,49^. This remains speculative at this point though. We currently do not know whether, and if so how, the length of the autocorrelation window in lower- and higher-order networks is related to different degrees of temporal integration of external and internal stimuli (see ^39^ for first steps in this direction).

### Limitations

Several limitations need to be mentioned. We employed a novel measure of time window, the ACW-0. While our findings show rest and task differences of ACW-0 compared to the shorter ACW-50, future studies both imaging and modelling may be necessary to further establish their differences. Yet another question in this respect is whether ACW-50 and ACW-0 mediate different cognitive processes as it is suggested by their association with different regions/networks in the spatial hierarchy, i.e., core and periphery. Modelling studies will be needed to establish a causal relation between rest and task temporo-spatial hierarchies. While employing different measures of rest-task similarity, we were unable to establish a causal rest-task connection in temporo-spatial hierarchies.

### Conclusion

We show the convergence of the brain’s temporal hierarchy, i.e., its intrinsic neural timescales, with the spatial topography of the core-periphery hierarchy during both rest and task states. This suggests an intrinsically temporospatial core-periphery hierarchy in the human cortex which shapes its cognitive processing during rest and task states. Pointing to its importance, the temporo-spatial nature of the brain’s core-periphery organization may be key in understanding mental features like self and consciousness including their temporo-spatial organization in perception ^27,32,33,50–52^. Specifically, the temporo-spatial core-periphery organization could provide a dynamic feature shared by both neural and mental levels, i.e., a “common currency” ^28^, requiring what recently has been described as “Spatiotemporal Neuroscience” ^53^.

## Methods

### Experimental model and subject details

The analyses involved magnetoencephalography (MEG) data of 89 subjects from the Human Connectome Project (HCP) WU-Minn HCP 1200 subjects data release ^54^. Resting state MEG data were acquired in runs of approximately 6 minutes. During the scan, the subjects were instructed to relax with eyes open and maintain fixation on a red crosshair. ECG and EOG recordings were also performed. Following the completion of resting state MEG, subjects were asked to complete three tasks of language processing (story vs. math, StoryM), motor (Motort) and working memory (Wrkmem). Task MEG data were acquired with the same parameters as the rest. Each task was approximately 7, 14 and 10 for StoryM, Motort and Wrkmem, respectively. The sampling frequency was 2034.5 Hz.

### Task paradigms

The tasks were all block-design paradigms. In the working memory task, participants were instructed to retain images in their visual working memory and compare them with subsequently presented images. The language processing task consisted of 7 blocks of a story task interleaved with 15 blocks of a math task. The story blocks presented participants with brief auditory stories. Motor processing was assessed using a task in which participants were presented with visual cues instructing the movement of either the right hand, left hand, right foot, or left foot. The paradigm of each task is extensively discussed in the WU-Minn HCP 1200 Subjects data release manual.

### Preprocessing

The HCP released preprocessed data. It included the following steps: removal of artifacts, bad channels, and bad segments based on HCP quality assurance standards ^55^, down-sample to 508.62 Hz, bandpass (1.3–150 Hz) and notch (59–61 Hz, 119–121 Hz) filtering, and removal of non-brain components through independent component analysis (ICA).

### Source reconstruction

was conducted similar to a recent article by Demirtaş et al.^56^, which used the same HCP data. Briefly, reconstruction was done for 8004 vertices (8k space) on the cortical surface using Fieldtrip and software provided by HCP. First, we applied a low-pass filter (50 Hz) on the sensor data, which was projected on to source space by synthetic aperture magnetometry. The details of the projection are provided in Demirtaş et al.^56^. After source reconstruction, time courses were parcellated using two well-known templates provided by Schaefer et al. ^40^ (Schaefer) and Ji et al. ^41^ (Ji). The Schaefer and Ji templates consist of 200 and 360 regions cortical areas, respectively. The regions Schaefer template were divided into 7 networks including Visual, Somatomotor, Dorsal Attention, Salience, Limbic, FPC (frontoparietal), and DMN (default-mode). Moreover, the Ji template consisted of 12 networks of Visual1, Visual2, Auditory, Somatomotor, Dorsal Attention, Posterior Multimodal, Ventral Multimodal, Orbito Affective, Language, Cingulo Opercular, FPC, and DMN. The templates were resampled to 8K space using resampling tools available in the HCP workbench software. Finally, the time courses were averaged over runs resulting in four conditions of Rest (resting state MEG), StoryM, Motort, and Wrkmem (task states MEG).

### Calculation of the metrics

In the next step, autocorrelation (ACF) of the regions’ time courses were calculated within the statsmodel library ^57^ using the fast Fourier transform algorithm. From that, two ACW values were extracted per autocorrelation function: the first lag (in milliseconds) where the autocorrelation decays to 50% of its maximum (ACW-50) and the first instance where the autocorrelation reaches zero (ACW-0).

### Core-periphery division

To analyze the core-periphery (CP) hypothesis, the regional data was divided into two categories of periphery and core base on the network they belonged to. Three different core-periphery divisions were defined. 1) Core-periphery based on the Schaefer template and the principle gradient in the Margulies et al. article^14^ (Schaefer/Margulies or SCP) in which the regions in the Visual, Somatomotor, Dorsal Attention and Salience networks were labelled as periphery and the regions in the Limbic, FPC, and DMN networks as the core. 2) Core-periphery based on the Ji template and the unimodal-transmodal definition in Ito et al.^1^ (Ji/Ito or JCP) in which periphery is defined as the regions in Visual1, Visual2, Auditory, and Somatomotor networks and core as the regions in Dorsal Attention, Posterior Multimodal, Ventral Multimodal, Orbito Affective, Language, Cingulo Opercular, FPC, and DMN. 3) Restricted core-periphery based on the Ji template and a restricted version of core from JCP definition (RCP) in which the periphery is similar to JCP, but the core is defined as only the regions in Cingulo Opercular, FPC, and DMN. This definition of CP ignores the rest of the regions.

### Anterior-posterior regression

The core-periphery organization of ACW was validated by removing the probable effect of the anterior-posterior organization from both ACW-50 and ACW-0. From the HCP released data, the averaged T1-weighted structural MRI was used to extract the y-coordinate of each region defined in both Schaefer and Ji templates. The ACW values for all regions and their corresponding y-coordinates were fed into a linear regression model as dependent and independent variables, respectively. The residual of the model was used in the CP analysis as the residual ACW.

### Rest-task similarity

The similarity between resting and task states was addressed using spatial correlation, linear regression and regional correlation. The spatial correlation was calculated as a single Pearson correlation coefficient between resting and a task condition (e.g. Rest vs. StoryM) over brain regions after averaging over subjects (thus creating a single brain per condition, illustrated in the box of Figure 4a). The spatial similarity was further investigated using linear regression. The regional ACW values were averaged over subjects. Then, task state ACW was modelled as a function of resting state for each task condition. The regional correlation was used to measure the regional similarity between resting and task states. This correlation was calculated for each region across subjects between a pair of conditions (illustrated in the box of Figure 4b).

### Rest-task difference

The difference between rest and task state was calculated as a percentage of change per region per subject. Each region’s rest and task (e.g. StoryM) values were put in the 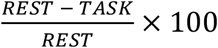 formula to *REST* both measures the change from the rest to the task and normalize against the rest at the same time. The percentage of change was averaged across subjects and used in the analysis of the core-periphery organization.

### Density estimation

To observe the relationship between ACW-50 and ACW-0, we estimated the probability distribution of the data, by removing all the labels (e.g. region and network), by averaging over regions, and by averaging over subjects. Removing all the labels means that for each scale (whether 50 or 0) all the data from all tasks, regions, and subjects were put together and fed to the estimation algorithm. Estimation was performed using the statsmodel’s kernel density estimator (KDEUnivariate) using the Gaussian kernel and expectation-maximization algorithm.

### Inter-individualeffects

To investigate the impact of inter-individual differences on ACW scales, we used logistic regression. A binary classification using logistic regression was designed. First, all ACW values were averaged over regions. Then, each subject was labelled either ‘high variability’ and ‘low variability’ (actual labels) based on their resting state ACW value. High variability refers to those subjects whose averaged ACW value (averaged over regions) is above the median of ACW values. Logistic regression was incorporated to determine the label for each subject during each task state. Then the predicted labels were compared to actual ones. The area under the Receiver Operating Characteristic (ROC) curve was computed based on the predicted and actual labels. ROC is a well-known metric for diagnostic test evaluation. In a ROC curve, the true positive rate is plotted as a function of the false positive rate for different cut-off points of a parameter (ACW-50 or ACW-0). The area under the ROC curve (AUROC) tells how much the model is capable of distinguishing between two diagnostic groups (low variability subjects compared to high variability ones). The higher the area under the curve, the better the model at correctly predicting the true label.

### Core-periphery Classification andfeature selection

To investigate which ACW scale is better at discriminating core from periphery regions, a 2-class classification problem was used with the logistic function as the classifier. Each region’s signal was labelled either core (class 1) or periphery (class 2). All the other labels were removed from the data (i.e. condition, subject, network, and region). For each region, the ACW values were used as features. Thus, each sample in the model had a 1-D feature vector containing either ACW-50 or ACW-0 value and a label (whether core or periphery). To increase the reliability of the results, we used the k-fold crossvalidation method with 20 folds. Three different models were created. Model 1 used both ACW values as features (each sample had two features), Model 2 used only ACW-50 and the third model used only ACW-0. The training and testing were conducted using the K-fold cross-validation algorithm with 20 folds. Accuracy, precision, recall, and the area under the AUC curve was calculated as efficiency metrics.

All of the classification steps were implemented using the scikit-learn library of Python programming language. For validation purposes, the same procedure was reused with a support vector machine (SVM) classifier using the radial basis function (RBF) kernel. Furthermore, mutual information ^43^ was used as a univariate feature selection method to determine which of the two scales is better if it was required to choose between the two. Mutual information is a data-driven method capable of capturing any kind of statistical dependency between the target class variable and the feature. It is equal to zero if the two are independent and the more they are dependent the more mutual information approaches 1. For that, we used the” SelectKBest” function with the “mutual_info_classif” scoring function from the scikit-learn library.

### Simulated Signals

The relationship between ACW-50 and ACW-0 was investigated using simulation analysis. 20000 pseudo-aleatory signals within four different categories (5000 each) were simulated. The autocorrelation function of each signal was calculated and from that ACW scales were extracted. The categories included (I) pink noise, (II) sine wave, and linear combinations of (III) pink noise and sine wave, and (IV) pink noise, white noise and sine wave. Uniformly distributed random weights were used for the linear combinations and the frequency of the sine waves. The frequency range and the sampling rate was fixed to be equal to our real data. Pink noise was chosen to model the scale-free behaviour, white noise for pure randomness and sine wave for oscillation. For each category two density functions were calculated (one for each ACW scale) and compared to the density functions of our real data.

### Statistics and Reproducibility

All statistical analyses were performed in the statsmodel library ^57^ and R v.3.6 and all p-values were corrected for multiple comparisons using the FDR correction method. Student’s t-test, One- and two-way analysis of variance (ANOVA) were used to determine the difference in ACW during resting and task states. For significant results, post-hoc analysis using Tukey’s HSD method was conducted to show the within factor differences. The steps for each analysis are described in their designated section both in results and method.

### Software

All steps of the data processing were performed with in-house scripts written in Python programming language using NumPy, SciPy, cifti, joblib, matplotlib, and seaborn libraries. The source code and the figures are freely available at www.georgnorthoff.com/codes/. For brain map visualization purposes, wb_view (part of Connectome Workbench software, https://www.humanconnectome.org/software/connectome-workbench) was used.

## Data Availability

The data that support the findings of this study are freely available at the Human Connectome Project’s repository at https://db.humanconnectome.org.

## Code Availability

The scripts that support the findings of this study are freely available at www.georgnorthoff.com/codes

## Acknowledgments

This research has received funding from the European Union’s Horizon 2020 Framework Program for Research and Innovation under the Specific Grant Agreement No. 785907 (Human Brain Project SGA2). GN is grateful for funding provided by UMRF, uOBMRI, CIHR and PSI. We are also grateful to Chris J. Honey for giving useful suggestions. We are also grateful to CIHR, NSERC, and SHERRC for supporting our tri-council grant from the Canada-UK Artificial Intelligence (AI) Initiative ‘The self as agent-environment nexus: crossing disciplinary boundaries to help human selves and anticipate artificial selves’ **(ES/T01279X/1) (together with Karl J. Friston from the UK)**

## Authors Contributions

Mehrshad Golesorkhi helped with study design, data analysis, simulation and manuscript writing. Javier Gomez-Pillar helped with method validation, simulation, and manuscript editing. Shankar Tumati helped with study design and manuscript editing. Maia Fraser helped with manuscript editing and idea validation. Georg Northoff as the principal investigator supervised the study and helped with study design, analysis guidance, manuscript writing, literature review and the submission process.

