## Supplementary Material for "Temporal hierarchy of intrinsic neural timescales converges with spatial core-periphery organization"

### Supplementary Figures

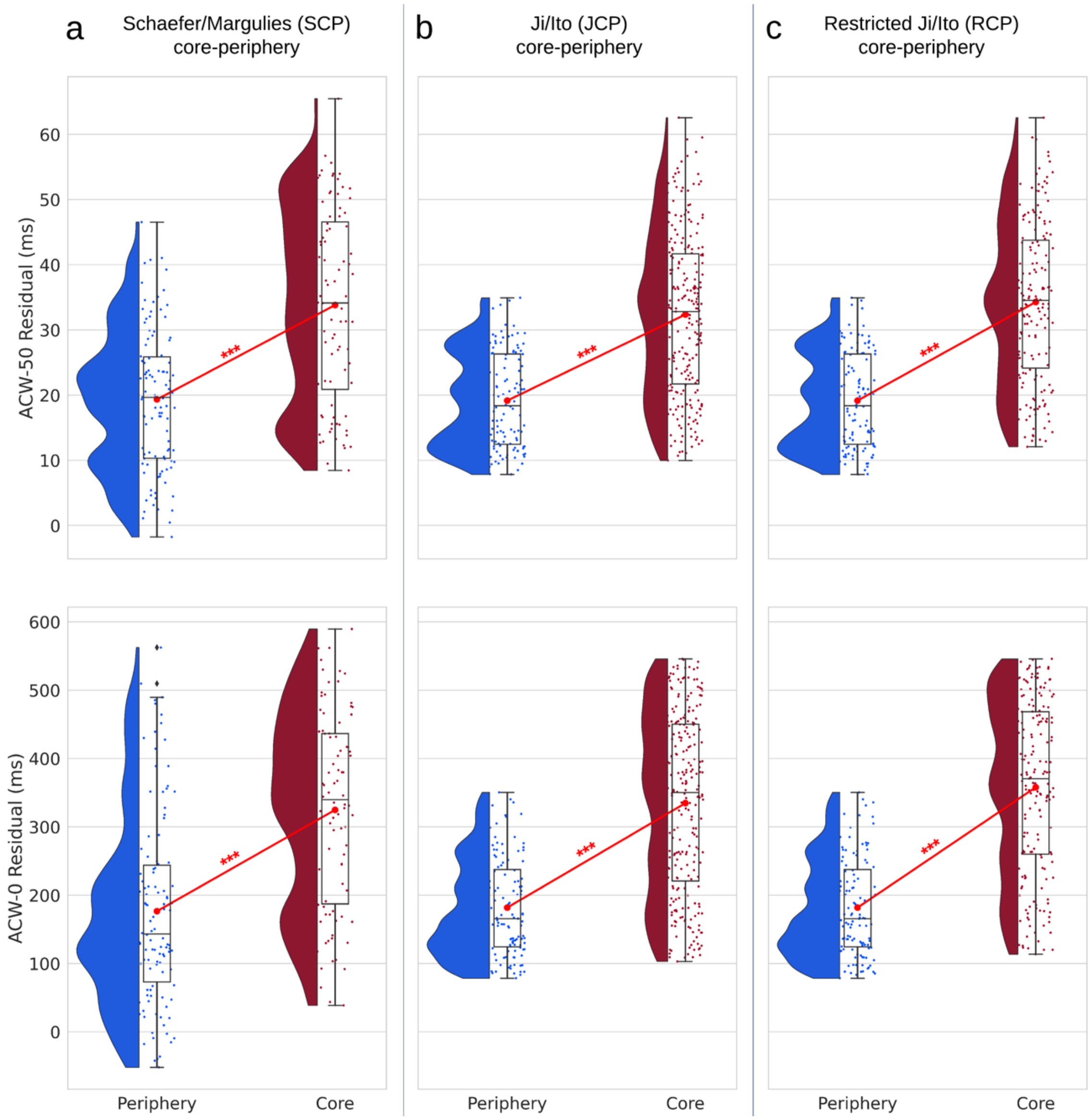


Supplementary Figure 1. Residual autocorrelation window (ACW) during resting state for the three core-periphery (CP) divisions after regressing out the anterior-posterior gradient. Rainclouds represent regions divided into core and periphery. Values are presented in milliseconds. The y-coordinate of each region's anatomical location was used as its index along the anterior-posterior axis. The y-coordinates were regressed out from ACW values in a linear regression model and then the residual ACW was plotted along the core-periphery division. The results replicate our original results with ACW during resting state. (a) Rainclouds for the ACW values of core and periphery using the Schaefer parcellation template (SCP): ACW-50, $t = -7.80, d = -1.14$, ACW-0, $t = -7.23, d = -1.04$ (b) The extended core definition of the Ji parcellation template (JCP): ACW-50, $t = -10.43, d = -1.20$, ACW-0, $t = -11.65, d = -1.33$ (c) The restricted definition of core (RCP): ACW-50, $t = -11.69, d = -1.43$, ACW-0, $t = -13.51, d = -1.62$. Stars represent the significance level ($*** \equiv \alpha = 0.001$).

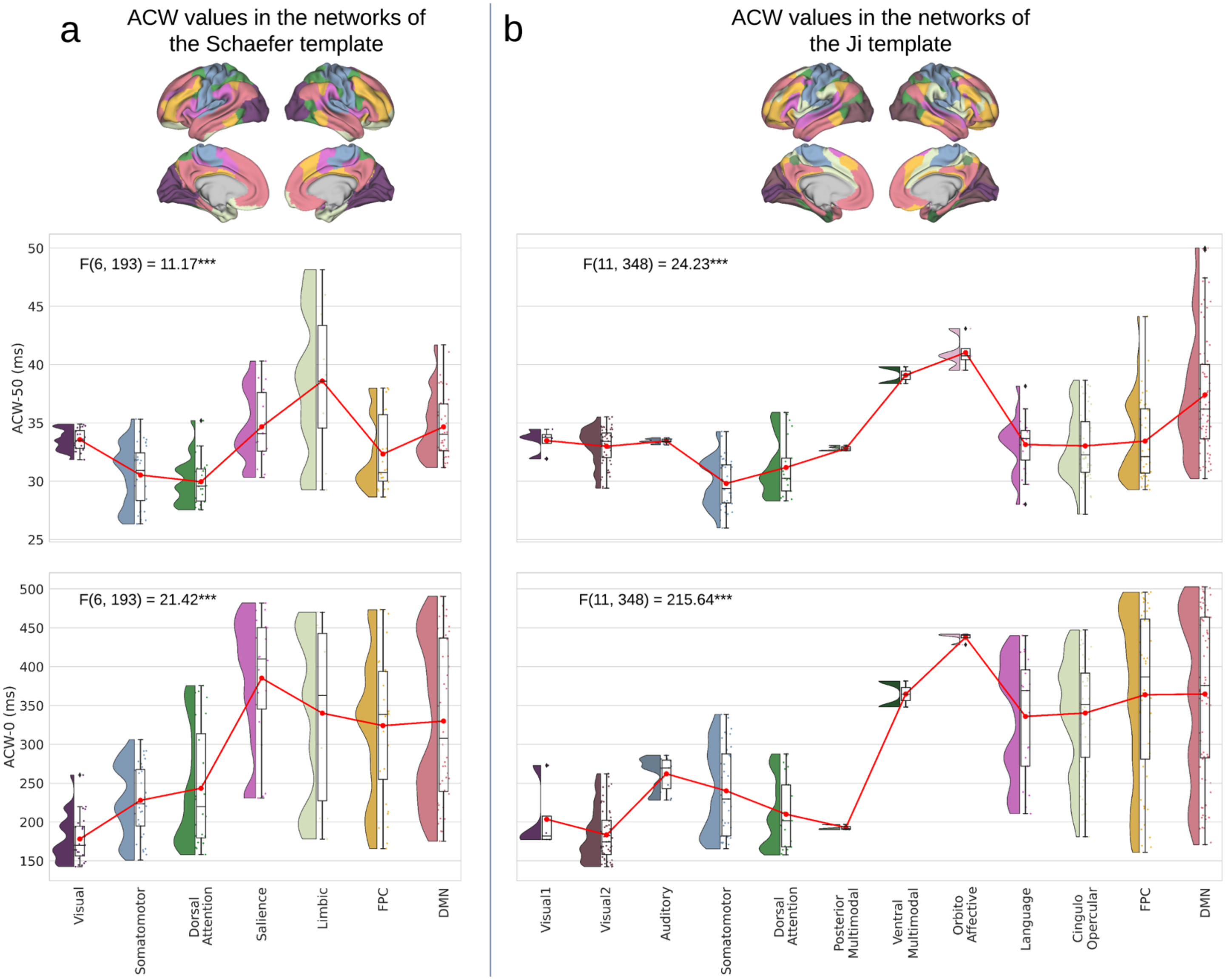


Supplementary Figure 2. Autocorrelation window (ACW) during resting state for the networks. Values are presented in milliseconds. Brain plots show network definitions in different templates. Rainclouds represent regions of different networks defined in (a) Schaefer and (b) Ji templates. Stars represent the significance level ($*** \equiv \alpha = 0.001$).

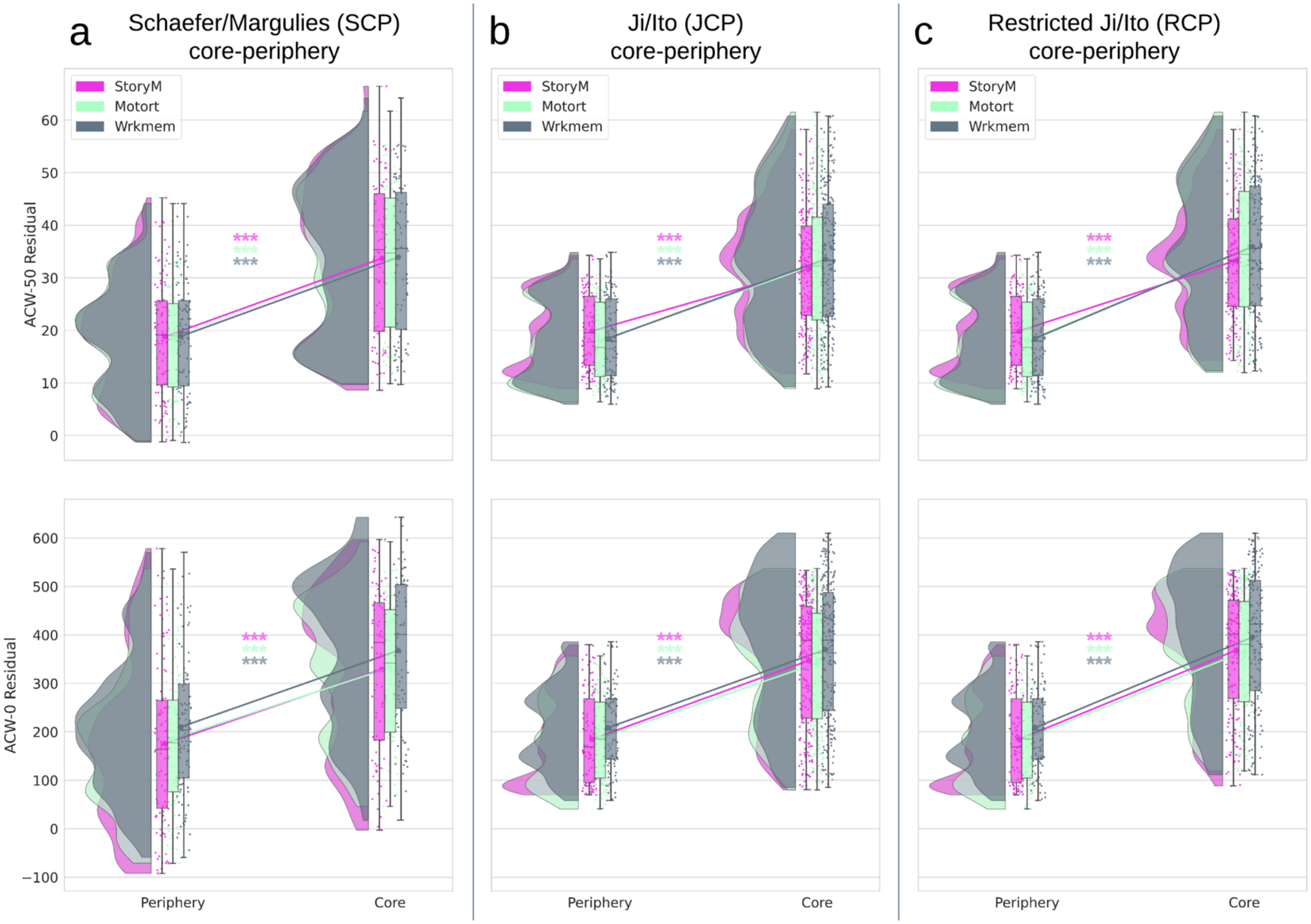


Supplementary Figure 3. Residual autocorrelation window (ACW) during different task states for the three core-periphery (CP) divisions after regressing out the anterior-posterior gradient. Rainclouds represent regions divided into core and periphery. The y-coordinates were regressed out from ACW values in a linear regression model and then the residual ACW was plot along the core-periphery division. The results replicate our original results for all tasks. (a) Rainclouds for the ACW values of core and periphery using the Schaefer parcellation template (SCP). The effect of core-periphery factor in the two-way ANOVA was significant for ACW-50 ($F(1,594) = 152.67, \eta^2 = 0.20$) and ACW-0 ($F(1,594) = 144.40, \eta^2 = 0.19$) (b) The extended core definition of the Ji parcellation template (JCP): ACW-50, $F(1,1074) = 514.64, \eta^2 = 0.32$; ACW-0, $F(1,1074) = 518.48, \eta^2 = 0.32$ (C) The restricted definition of core (RCP): ACW-50, $F(1,1074) = 565.43, \eta^2 = 0.38$ ; ACW-0, $F(1,1074) = 607.97, \eta^2 = 0.40$. Stars represent the significance level ($*** \equiv \alpha = 0.001$).

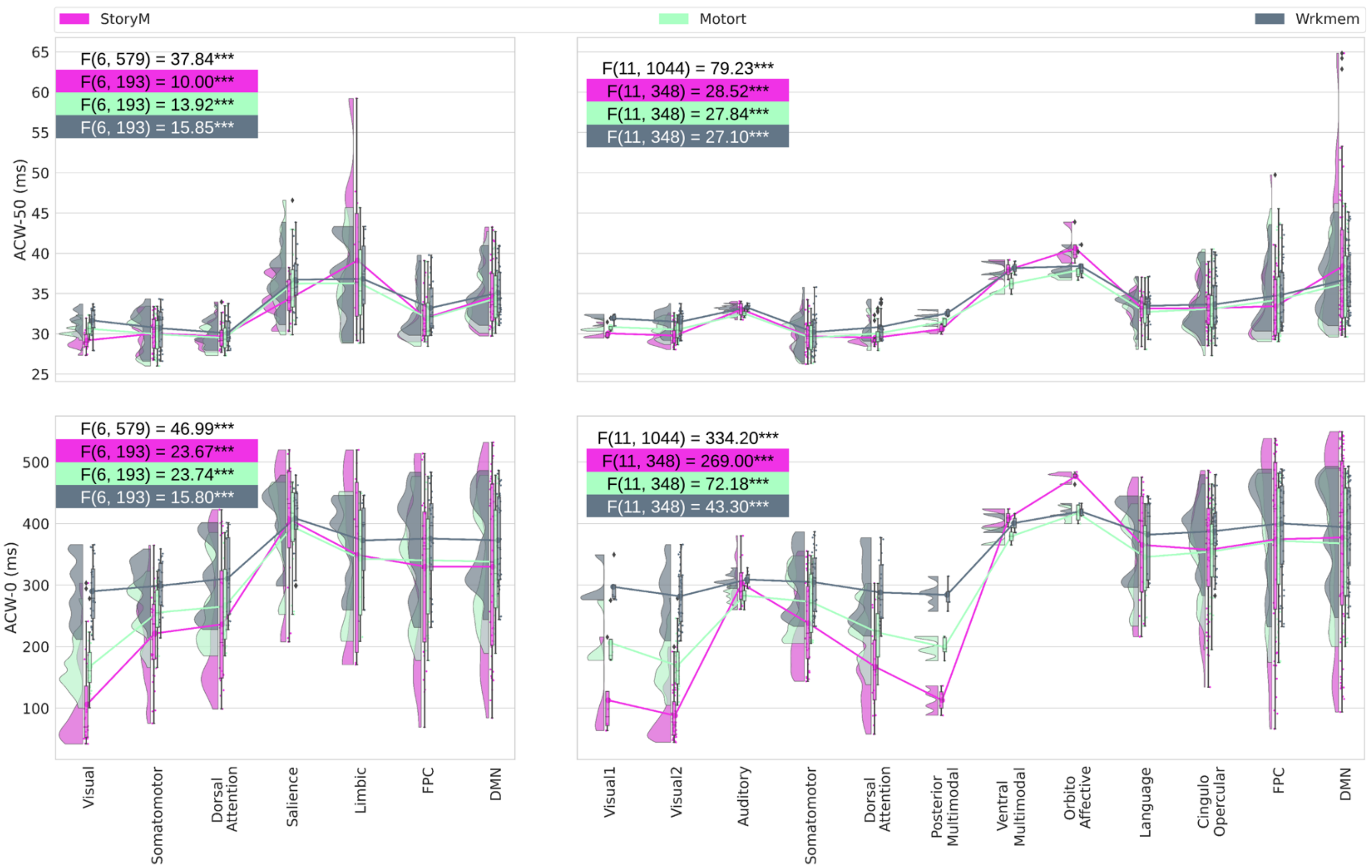


Supplementary Figure 4. Autocorrelation window (ACW) during different task states for the networks. Values are presented in milliseconds. Rainclouds represent regions of different networks defined in Schaefer (left) and Ji (right) templates. Stars represent the significance level ($*** \equiv \alpha = 0.001$). For each combination of template and scale (e.g. Schaefer and ACW-50, top left), first, a two-way ANOVA was calculated using task and network as factors to determine if network has any significant effect over the whole data. Then, a one-way ANOVA was calculated for each task using only the network as the factor to investigate the effect of network on each task. Values are presented in Supp. Table 2.

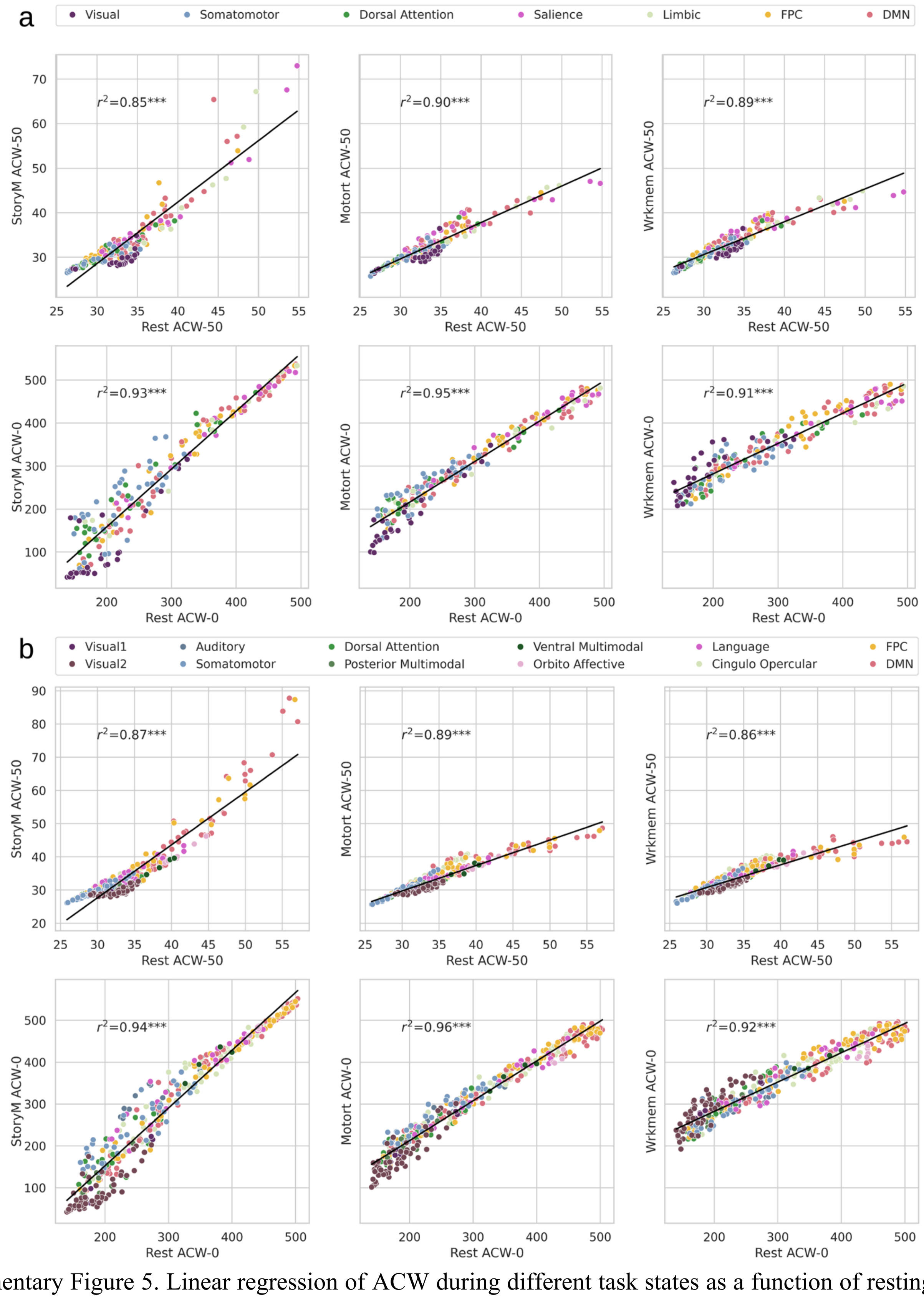


Supplementary Figure 5. Linear regression of ACW during different task states as a function of resting state for the Schaefer (a) and Ji (b) parcellation templates. Each scatter plot represents a task as a function of the resting state. Each dot represents a region's ACW value averaged over subjects. The linear relationship in all conditions is significant at $\alpha = 0.001$ and they indicate that at least 85% variance of the task is explained by resting state.

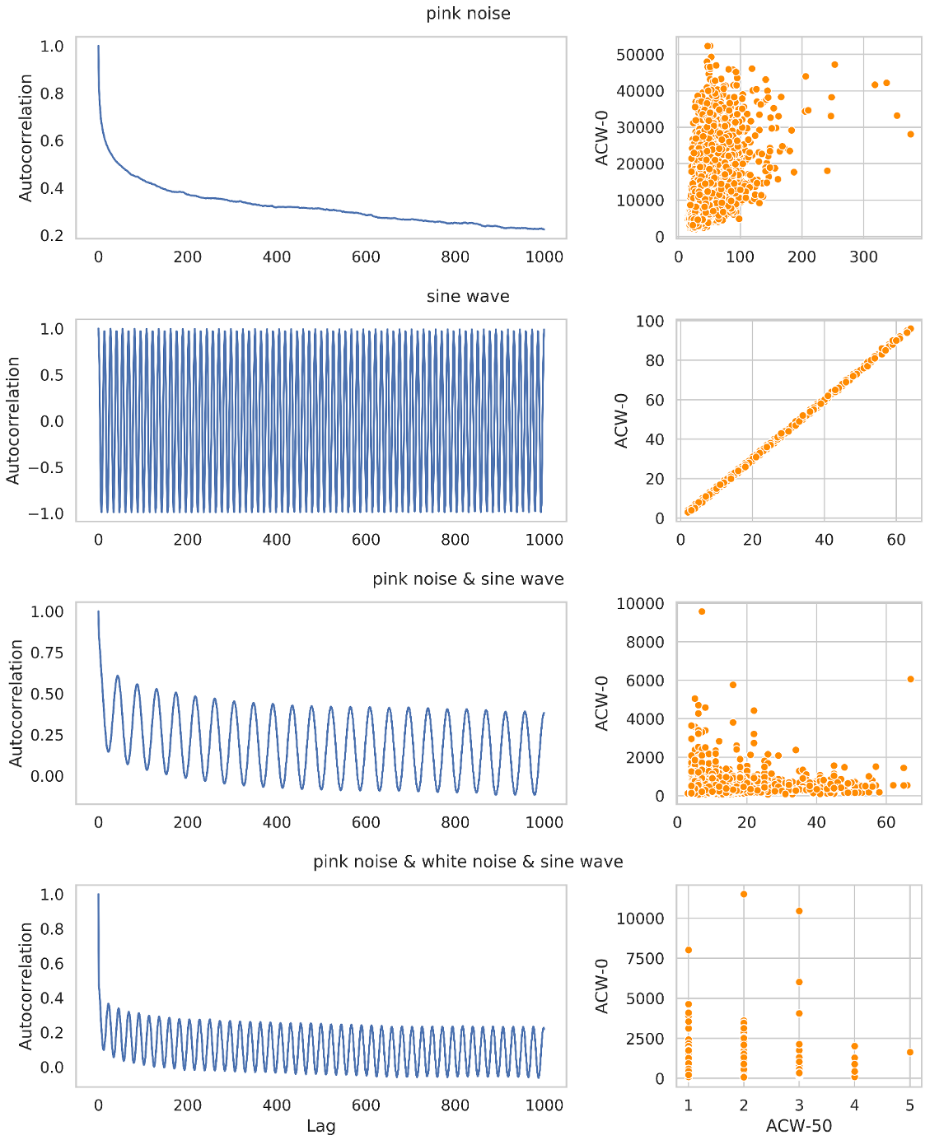


Supplementary Figure 6. ACW scales on simulated signals. Left shows the autocorrelation function (first 1000 lags) of a sample for each category of the simulated signals. Right shows the scatter plots for ACW-50 and ACW-0 of all signals in each category.

### Supplementary Tables

Supplementary Table 1. Two-way ANOVA analysis for different core-periphery organizations during task state. Task condition (3 levels) and core-periphery (CP, 2 levels) where used as independent factors.

| | | | |
|---|---|---|---|
| SCP | ACW-50 | Task | $F(2{,}594) = 1.93, \eta^2 = 0.005, p = 0.15$ |
| | | CP | $F(1{,}594) = 57.69, \eta^2 = 0.087, p < 0.001$ |
| | | Interaction | $F(2{,}594) = 0.76, \eta^2 = 0.002, \mathrm{p} = 0.46$ |
| | ACW-0 | Task | $F(2{,}594) = 26.13, \eta^2 = 0.072, p < 0.001$ |
| | | CP | $F(1{,}594) = 70.47, \eta^2 = 0.097, p < 0.001$ |
| | | Interaction | $F(2{,}594) = 2.21, \eta^2 = 0.006, \mathrm{p} = 0.10$ |
| JCP | ACW-50 | Task | $F(2{,}1074) = 7.51, \eta^2 = 0.010, p < 0.001$ |
| | | CP | $F(1{,}1074) = 318.44, \eta^2 = 0.223, p < 0.001$ |
| | | Interaction | $F(2{,}1074) = 7.62, \eta^2 = 0.010, p < 0.001$ |
| | ACW-0 | Task | $F(2{,}1074) = 77.24, \eta^2 = 0.089, p < 0.001$ |
| | | CP | $F(1{,}1074) = 464.40, \eta^2 = 0.267, p < 0.001$ |
| | | Interaction | $F(2{,}1074) = 20.59, \eta^2 = 0.023, p < 0.001$ |
| RCP | ACW-50 | Task | $F(2{,}885) = 7.26, \eta^2 = 0.011, p < 0.001$ |
| | | CP | $F(1{,}885) = 311.75, \eta^2 = 0.254, p < 0.001$ |
| | | Interaction | $F(2{,}885) = 7.39, \eta^2 = 0.012, p < 0.001$ |
| | ACW-0 | Task | $F(2{,}885) = 69.47, \eta^2 = 0.085, p < 0.001$ |
| | | CP | $F(1{,}885) = 549.90, \eta^2 = 0.339, p < 0.001$ |
| | | Interaction | $F(2{,}885) = 23.14, \eta^2 = 0.028, p < 0.001$ |

Supplementary Table 2. Two-way ANOVA analysis for different core-periphery organizations during task state for residual ACW. Task condition (3 levels) and core-periphery (CP, 2 levels) where used as independent factors.

| | | | |
|---|---|---|---|
| SCP | ACW-50 | Task | $F(2{,}594) = 0.24, \eta^2 = 0.006, p < 0.001$ |
| | | CP | $F(1{,}594) = 152.67, \eta^2 = 0.204, p < 0.001$ |
| | | Interaction | $F(2{,}594) = 0.12, \eta^2 = 0.003, p < 0.001$ |
| | ACW-0 | Task | $F(2{,}594) = 2.98, \eta^2 = 0.008, p < 0.001$ |
| | | CP | $F(1{,}594) = 144.40, \eta^2 = 0.193, p < 0.001$ |
| | | Interaction | $F(2{,}594) = 0.03, \eta^2 = 0.0009, p < 0.001$ |
| JCP | ACW-50 | Task | $F(2{,}1074) = 3.31, \eta^2 = 0.004, p < 0.001$ |
| | | CP | $F(1{,}1074) = 514.64, \eta^2 = 0.322, p < 0.001$ |
| | | Interaction | $F(2{,}1074) = 0.19, \eta^2 = 0.0002, p < 0.001$ |
| | ACW-0 | Task | $F(2{,}1074) = 5.11, \eta^2 = 0.006, p < 0.001$ |
| | | CP | $F(1{,}1074) = 518.46, \eta^2 = 0.323, p < 0.001$ |
| | | Interaction | $F(2{,}1074) = 0.10, \eta^2 = 0.0001, p < 0.001$ |
| RCP | ACW-50 | Task | $F(2{,}885) = 2.99, \eta^2 = 0.004, p < 0.001$ |
| | | CP | $F(1{,}885) = 565.43, \eta^2 = 0.388, p < 0.001$ |
| | | Interaction | $F(2{,}885) = 0.26, \eta^2 = 0.0003, p < 0.001$ |
| | ACW-0 | Task | $F(2{,}885) = 5.05, \eta^2 = 0.006, p < 0.001$ |
| | | CP | $F(1{,}885) = 607.97, \eta^2 = 0.404, p < 0.001$ |
| | | Interaction | $F(2{,}885) = 0.10, \eta^2 = 0.0001, p < 0.001$ |

Supplementary Table 3. Post-hoc results for periphery vs. core in different task conditions for residual ACW. Tukey HSD method was used to determine the between factor significance of ANOVA on both residual ACW-50 and residual ACW-0 along different CP divisions.

| | | StoryM | | | Motort | | | Wrkmem | | |
|---|---|---|---|---|---|---|---|---|---|---|
| | | $t$ | $d$ | $p$ | $t$ | $d$ | $p$ | $t$ | $d$ | $p$ |
| SCP | ACW-50 | -7.72 | -1.10 | *** | -6.99 | -0.99 | *** | -7.21 | -1.02 | *** |
| | ACW-0 | -6.88 | -0.98 | *** | -6.74 | -0.96 | *** | -7.10 | -1.01 | *** |
| JCP | ACW-50 | -11.04 | -1.25 | *** | -10.31 | -1.16 | *** | -10.59 | -1.20 | *** |
| | ACW-0 | -11.67 | -1.32 | *** | -11.17 | -1.26 | *** | -11.63 | -1.31 | *** |
| RCP | ACW-50 | -12.82 | -1.53 | *** | -11.88 | -1.41 | *** | -12.16 | -1.45 | *** |
| | ACW-0 | -12.97 | -1.54 | *** | -12.68 | -1.51 | *** | -13.27 | -1.58 | *** |
| *d stands for Cohen's d effect size. The values are represented as periphery vs. core.* *Stars represent the significance level (*** ≡ $\alpha = 0.001$, ** ≡ $\alpha = 0.01$).* | | | | | | | | | | |

Supplementary Table 4. Two-way and one-way ANOVA analysis for networks in different ACW task states. First, a two-way ANOVA (task and network as the independent factors) was calculated to determine the overall effect of network on ACW scales, and then for each task, a one-way ANOVA was calculated using network as the independent factor.

| | | | |
|---|---|---|---|
| Schaefer template | ACW-50 | Overall | $F(6, 579) = 37.48, \eta^2 = 0.27$ |
| | | StoryM | $F(6, 193) = 10.00, \eta^2 = 0.23$ |
| | | Motort | $F(6, 193) = 13.92, \eta^2 = 0.30$ |
| | | Wrkmem | $F(6, 193) = 15.85, \eta^2 = 0.33$ |
| | ACW-0 | Overall | $F(6, 579) = 46.99, \eta^2 = 0.29$ |
| | | StoryM | $F(6, 193) = 23.67, \eta^2 = 0.42$ |
| | | Motort | $F(6, 193) = 23.74, \eta^2 = 0.42$ |
| | | Wrkmem | $F(6, 193) = 18.80, \eta^2 = 0.32$ |
| Ji template | ACW-50 | Overall | $F(11, 1044) = 79.23, \eta^2 = 0.43$ |
| | | StoryM | $F(11, 348) = 28.52, \eta^2 = 0.47$ |
| | | Motort | $F(11, 348) = 27.84, \eta^2 = 0.46$ |
| | | Wrkmem | $F(11, 348) = 27.10, \eta^2 = 0.46$ |
| | ACW-0 | Overall | $F(11, 1044) = 334.20, \eta^2 = 0.67$ |
| | | StoryM | $F(11, 348) = 269.00, \eta^2 = 0.89$ |
| | | Motort | $F(11, 348) = 72.18, \eta^2 = 0.69$ |
| | | Wrkmem | $F(11, 348) = 43.30, \eta^2 = 0.57$ |

Supplementary Table 5. Two-way ANOVA analysis for different core-periphery organizations in the change from resting to task state. Task condition (3 levels) and core-periphery (CP, 2 levels) where used as independent factors.

| | | | |
|---|---|---|---|
| SCP | ACW-50 | Task | $F(2{,}594) = 23.11, \eta^2 = 0.069, p < 0.001$ |
| | | CP | $F(1{,}594) = 10.63, \eta^2 = 0.016, p < 0.001$ |
| | | Interaction | $F(2{,}594) = 4.77, \eta^2 = 0.014, p < 0.001$ |
| | ACW-0 | Task | $F(2{,}594) = 71.38, \eta^2 = 0.183, p < 0.001$ |
| | | CP | $F(1{,}594) = 12.17, \eta^2 = 0.015, p < 0.001$ |
| | | Interaction | $F(2{,}594) = 14.80, \eta^2 = 0.038, p < 0.001$ |
| JCP | ACW-50 | Task | $F(2{,}1074) = 45.48, \eta^2 = 0.069, p < 0.001$ |
| | | CP | $F(1{,}1074) = 99.61, \eta^2 = 0.075, p < 0.001$ |

| | | | |
|---|---|---|---|
| | | Interaction | $F(2,1074) = 25.42, \eta^2 = 0.038, p < 0.001$ |
| | ACW-0 | Task | $F(2,1074) = 161.51, \eta^2 = 0.208, p < 0.001$ |
| | | CP | $F(1,1074) = 26.74, \eta^2 = 0.017, p < 0.001$ |
| | | Interaction | $F(2,1074) = 63.58, \eta^2 = 0.081, p < 0.001$ |
| RCP | ACW-50 | Task | $F(2,885) = 36.77\ \eta^2 = 0.066, p < 0.001$ |
| | | CP | $F(1,885) = 97.35, \eta^2 = 0.088, p < 0.001$ |
| | | Interaction | $F(2,885) = 25.09, \eta^2 = 0.045, p < 0.001$ |
| | ACW-0 | Task | $F(2,885) = 123.30, \eta^2 = 0.187, p < 0.001$ |
| | | CP | $F(1,885) = 34.92, \eta^2 = 0.026, p < 0.001$ |
| | | Interaction | $F(2,885) = 75.56, \eta^2 = 0.114, p < 0.001$ |
